# Functional characterization of the 9q34.13 locus identifies *RAPGEF1* as modulating risk for melanoma and nevi via RAS activation

**DOI:** 10.64898/2025.12.03.691936

**Authors:** Rohit Thakur, Mai Xu, Alexandra M. Thornock, Joshuah Yon, Samuel Anyaso-Samuel, Martin Lauss, Thomas Rehling, Linh Bui-Raborn, Hayley Sowards, Gerard Duncan, Lea Jessop, Timothy Myers, Raj Chari, Erping Long, Karen Funderburk, Jinhu Yin, Rebecca Hennessey, Emory Hseih, Hannah Levin, Mitchell J. Machiela, Tongwu Zhang, Goran Jonsson, D Timothy Bishop, Julia Newton-Bishop, Jeremie Nsengimana, Mark M. Iles, Maria Teresa Landi, Matthew H. Law, Thorkell Andresson, Jiyeon Choi, Leonard I. Zon, Jianxin Shi, Melanoma Meta-Analysis Consortium, Kevin M. Brown

## Abstract

Genome-wide association studies identified a melanoma- and nevus count-associated locus on chromosome band 9q34.13. Fine-mapping and melanocyte expression data collectively suggest two potential causal genes with opposite association with risk: higher levels of Rap guanine nucleotide exchange factor 1 (*RAPGEF1*) and lower levels of uridine-cytidine kinase 1 (*UCK1*). Colocalization analyses and conditional TWAS suggest multiple causal *cis*-regulatory sequence variants in partial linkage disequilibrium (LD) to each other. Melanocyte capture-HiC and CRISPR-inhibition demonstrated regulatory interactions between fine-mapped variants and the *RAPGEF1* and *UCK1* promoters. Focusing on *RAPGEF1*, we demonstrate *RAPGEF1* expression promotes melanocyte growth and drives malignant transformation of human immortalized melanocytes. Following treatment with human EGF, *RAPGEF1* overexpression activated both RAP1 and RAS. Further, we show *RAPGEF1* expression is significantly enriched in melanomas lacking strongly activating RAS-MAPK mutations, suggesting that *RAPGEF1* may promote oncogenic RAS-MAPK signaling in melanomas. Furthermore, in these tumors, we provide preliminary evidence to support the prognostic relevance of *RAPGEF1* expression in patients lacking *RAS* or *BRAF* mutations. Together with other recent studies, these data suggest that germline variation influencing RAS activation may play a key role in nevus development and melanoma risk.

## Introduction

Meta-analysis of genome-wide association studies (GWAS) for melanoma have identified 54 loci and 68 independent signals associated with melanoma risk [1]. Despite the success in identifying loci, the causal sequence variants and specific genes at these loci that influence risk and melanoma development remain to be established. Linkage disequilibrium (LD) typically results in numerous risk-associated variants, complicating efforts to identify causal variants. Further, most fine-mapped variants at complex trait loci identified by GWAS are not protein coding; instead most causal variants alter *cis*-regulatory activity of regulatory sequences including enhancers and promoters [2, 3]. Given that enhancers can act at distances of 1Mb or more [4, 5], numerous genes near risk-associated loci may be plausible causal candidates.

Identifying causal sequence variants and their gene targets is important to understanding potentially unappreciated key processes underlying risk, tumor development, and or tumor progression. Of known risk loci, the majority are also associated with heritable melanoma risk-associated traits [1], including hair, skin or eye color [1, 6], telomere length [1, 7–9], and number of melanocytic nevi [1, 10–12]. While these pleiotropic associations can provide clues to the likely causal genes at melanoma risk loci, establishing causal genes for most loci remains a challenge.

We and others have previously generated multiple genome- or GWAS-scale genomic, transcriptomic, epigenomic, chromatin conformation, and functional screening datasets in melanocytic cells or tissues relevant to melanoma risk [13–24]. Here we layer analyses of these datasets and characterize a melanoma risk and nevus count locus on chromosome band 9q34.13 [1] near the 31-end of RAP guanine nucleotide exchange factor 1 (*RAPGEF1*, also known as C3G) and establish the *RAPGEF1* gene as one of two potential causal candidate genes at this locus. We demonstrate that *RAPGEF1* expression promotes melanocyte proliferation, increases colony formation in immortalized melanocytes, and leads to both RAP1 and RAS activation. Finally, we demonstrate higher expression of *RAPGEF1* in melanoma tumors lacking RAS-pathway driver mutations consistently across multiple tumor sequencing datasets. These data suggest that germline modulation of RAS activity may play a key role in both nevus and melanoma development.

## Methods

### Genetic fine-mapping at the melanoma risk locus on chromosome band 9q34.13

We fine-mapped the melanoma risk locus on chromosome band 9q34.13 (a ±500 Kb window surrounding rs3780269) to identify potential causal variants. We applied multiple Bayesian fine-mapping methods including SuSiE (Sum of Single Effects) [25, 26], DAP-G [27, 28] (Deterministic Approximation of the Posteriors), and RSparsePro [29], which is robust to mismatching LD between the GWAS and the LD-reference panels. We used default parameters for each method. The min_abs_corr parameter was set to 0 for SuSiE. Melanoma GWAS Z-score values for each variant were from the fixed-effect inverse variance weighted meta-analysis of the full set of confirmed and self-reported melanoma cases and control, as described previously by Landi and colleagues [1]. We used a pre-computed LD matrix of 5,000 UK Biobank individuals as described previously [1]. While we allowed for a maximum of five credible sets during fine-mapping, we only retained individual credible sets where the cumulative posterior inclusion probability was > 0.85, and retained all variants in the credible set where the individual variant posterior inclusion probability was > 0.005. To inclusively fine-map variants, we also independently selected any variants with log likelihood ratio (LLR 1:1000) relative to the GWAS lead variant rs3780269. Finally, to assess variants that were not successfully genotyped or imputed in the GWAS (including notably insertion/deletion variants that were not a part of the Haplotype Reference Consortium imputation panel), we included any such variants that had an LD r^2^ > 0.8 (1000 Genomes Project, Phase 3, Version 5, EUR population) [30] with rs3780269 as identified by the LDlinkR package [31–33].

To further assess the possibility of other potential independent signals in the region, we conditioned on the lead GWAS variant (rs3780269) using the Genome-wide Complex Trait Analysis (GCTA, v1.94.1) [34] COnditional and JOint association (COJO) module [35], using default settings, a genomic window of ±1 Mb from the lead variant, and the UKBB LD matrix described above.

### Annotation of fine-mapped variants

We assessed all fine-mapped variants at this locus for overlap with melanocytic regulatory elements (defined by chromatin states, or accessibility) using the BedtoolsR package (intersect function) [36, 37] and epigenomic datasets from melanocytes and melanoma cell line models. We classified genomic regions as enhancers or promoters using the Roadmap project chromHMM data from two primary melanocyte cultures [17, 20]. Using this data, enhancers were defined using the following chromHMM states from all chromHMM tracks (primary: Enh, EnhG, EnhBiv; auxiliary: EnhG1, EnhG2, EnhA1, EnhA2, EnhWk, EnhBiv; and imputed: TxEnh5, TxEnh3, TxEnhW, EnhA1, EnhA2, EnhAF, EnhW1, EnhW2, EnhAc). Melanoma-specific enhancer annotations included 4_EnhA, 5_EnhM, 6_EnhW, 7_TxEnhM, 7_TxEnhW, and 9_TxWkEnhW, from a tumorigenic melanoma cell model [21]. Promoter states were defined in melanocytes from imputed ChromHMM as PromU, PromD1, PromD2, TssA, PromP, PromBiv, and Tx_Reg; and in melanoma cells as 1_TssA, 2_PromWkD, and 3_TssWkP. We analyzed our previously generated ATAC-seq data from five independent melanocyte cultures [22], along with publicly available omni-ATAC-seq data from nine melanoma cell lines derived from patient biopsies [19]. Any region annotated as enhancer or promoter or that overlapped ATAC-peak in either melanocyte or melanoma cells was considered *cis*-regulatory.

Allele-specific transcriptional activity of these variants was previously assessed using Massively Parallel Reporter Assays (MPRA) in immortalized primary melanocytes and UACC903 melanoma cells [23]. All fine-mapped variants for the melanoma risk signal were included in the MPRA library design, however one failed data QC and was subsequently excluded from analysis. Allelic impacts on transcriptional activity were assessed using a linear regression model, accounting for strand and transfection batch as covariates. A scrambled 145-bp sequence was used as a negative control and as a baseline to assess relative enhancer activity of specific regions surrounding fine-mapped variants. Allele-specific differences for each variant were tested relative to scrambled controls. Multiple hypothesis testing was corrected using Benjamini-Hochberg false discovery rate.

We assessed the impact of fine-mapped variants on the protein-coding sequence Variant effect predictor tool (https://grch37.ensembl.org/Homo_sapiens/Tools/VEP) [38] using GENCODE version 19 protein-coding transcripts.

### Cell culture

Frozen aliquots of melanocytes isolated from foreskin of healthy newborn males, mainly of European descent, were obtained from the Specialized Programs of Research Excellence (SPORE) in Skin Cancer Specimen Resource Core at Yale University following an established protocol [39]. Melanocytes were grown in Dermal Cell Basal Medium (American Type Culture Collection/ATCC PCS-200-030) supplemented with Melanocyte Growth Kit (ATCC PCS-200-041) and 1% amphotericin B/penicillin/streptomycin (120-096-711, Quality Biological) at 37°C with 5% CO_2_. Immortalized C283T [18] melanocytes as well as C283T cells stably expressing dCas9-KRAB [22] were generated as previously described and grown in identical conditions to primary melanocytes. All cells tested negative for mycoplasma contamination via MycoAlert PLUS Mycoplasma Detection Kit (LT07-710, Lonza).

### Assessing micro-RNA binding for variant overlapping RAPGEF1 3’-UTR region

To assess whether melanoma GWAS risk variant rs3739497, which resides in the *RAPGEF1* 31 untranslated region (UTR), is involved in potential post-transcriptional regulation of *RAPGEF1* through micro-RNAs (miRNAs), we assessed predicted miRNA binding sites near this variant. We queried several databases, including TargetScan [40], miRTarBase [41], miRDB [42, 43], and mirCode [44]. Binding sites that overlapped or were located within ±50 nucleotides of the variant position were considered candidate variant-dependent miRNA sites. Finally, to explore post-transcriptional regulation via sequence elements other than miRNAs, we queried the 3’UTR variant region against the AREsite database [45] that provides a genome-wide catalogue of AU-rich elements.

### Small RNA sequencing data from primary melanocytes and melanomas

To assess small RNA levels in primary human melanocytes (n=106), total RNA, including microRNA (miRNA), was purified using Qiagen miRNeasy kit (#217004). Briefly, cells were lyzed in Trizol and heated at 65C followed by chloroform extraction. The aqueous phase was further purified using a miRNeasy column following manufacturer’s instruction. miRNA sequencing libraries were constructed using NEB Next Small RNA Library Prep kit (E7330s) and the sequencing was performed on HiSeq2500. All samples show the proportion of Q30 or higher bases of at least 93%. 28-79 million reads were achieved for each sample with overall alignment percentage between 67-85%. Sequencing reads were processed using the mirMaster pipeline [46]. Quality control stage processing steps included removal of 3⍰ adapter sequences (AGATCGGAAGAGCACACGTCTGAACTCCAGTCAC), trimming of low-quality bases, and retention of reads that exceeded the default minimum length thresholds. Reads were then aligned to the human genome (GRCh38) using Bowtie [47], allowing up to five genomic alignments per read. miRNAs were quantified by aligning sequencing reads against the mature and precursor miRNA reference sequences from miRBase v22.1 [48–53]. Alignments permitted up to one mismatch. Expression values were normalized across samples using quantile normalization. For each miRNA, percentile rank of expression across the dataset was calculated based on median expression values.

*RAPGEF1* mRNA expression levels were available from the bulk RNA-seq data we previously simultaneously generated from the same 106 melanocyte cultures [13]. The correlation of *RAPGEF1* mRNA expression levels with candidate miRNAs was measured using Spearman’s rank correlation.

Expression levels of micro-RNAs in TCGA melanoma samples [14] were accessed through FirebrowseR (https://github.com/mxxdxxx/FirebrowseR).

### Quantitative Trait Locus (QTL) analysis of melanocytic cells

We assessed FDR significant QTL signals near the GWAS risk locus using previously-published melanocyte and melanoma datasets [9, 12]. Melanocyte expression QTL (eQTL), splice QTL (sQTL), and methylation QTL (meQTL) datasets were generated from 106 primary melanocyte cultures derived from individuals of European descent. Methylation probes from meQTL data were assigned to genes based on CpG location within 1.5 kb of the TSS, 5⍰-UTR, 1st exon, gene body, or 3⍰-UTR of a gene. Tumor eQTL data were derived from TCGA melanoma primary tumors as a part of the PancanQTLv2.0 eQTL study; only data from nominally significant SNP-gene pairs (P<0.01) were made available [79] (https://hanlaboratory.com/PancanQTLv2/).

GWAS colocalization using the melanocyte eQTL dataset was performed using the SuSiE + Coloc [54] and SharePro tools [55]. SuSiE + Coloc performs fine-mapping of GWAS and QTL signals via SuSiE, followed by colocalization via the Coloc tool. SharePro leverages linkage disequilibrium (LD) to cluster correlated variants into effect groups, followed by colocalization. GWAS summary data, and the *RAPGEF1* and *UCK1* melanocyte eQTL summary data (± 500 Kb around the lead variant) were used as input for this analysis. Default parameters were used except the sigma parameter, which represents the prior probability of colocalization between traits. Since the region being assessed harbors multiple GWAS/QTL signals with complex LD, we provide results using multiple sigma values, ranging from most stringent (1 x 10^-5^) to less stringent (1 x 10^-4^, 1 x 10^-3^). However given that relaxing this value from the default of 1 x 10^-5^ increases the potential for false positive findings, we formally tested colocalization using this default value. Colocalization probability > 0.8 was considered strong evidence supporting colocalization, while a colocalization probability < 0.2 was considered evidence against colocalization [55].

We also used these QTL datasets to perform transcriptome (melanocyte and TCGA melanoma tumors) and methylome-wide (melanocyte) association studies (TWAS/MWAS). To identify conditionally independent loci, we performed conditional TWAS analysis using the FUSION tool [56].

### Capture-HiC assay for linking risk variants to target genes

As described previously [22], we performed melanoma GWAS-region focused capture Hi-C assays in primary human melanocytes to facilitate variant-to-gene mapping across all 68 melanoma genome-wide significant signals. Capture probes were designed by Arima Genomics (San Diego, CA) using 2x tiling with least stringent masking and XTHSBoosting, resulting in an Agilent SureSelect library (Santa Clara, CA) targeting all restriction fragments generated by DpnII and MboI (recognition sites: GATC, GANTC) within each association interval. We generated 15 capture Hi-C libraries from five independent primary melanocyte cultures (C56, C140, C205, C24, and C27), with three technical replicates prepared per culture. Barcoded libraries were pooled and sequenced on an Illumina NovaSeq platform across one SP and one S1 flow cell, yielding ~5.7 billion 150 bp paired-end reads in total.

Sequencing reads were processed using the HiCUP pipeline [57] with alignment to the hg19 reference genome via Bowtie2 [58]. Technical replicates from each melanocyte culture were combined prior to downstream analyses. Chromatin interactions were detected using the CHiCAGO pipeline (v1.16.0) [59] at both individual restriction fragment-level and aggregated four-fragment resolution, as previously described [22]. Interactions with CHiCAGO scores >5 were considered high-confidence visualized on the WashU [60, 61] and UCSC genome browsers [62–66]. To mitigate cross-linking variability and restriction fragment boundary effects, we expanded each fine-mapped variant into a ±500 bp window and evaluated interactions from both the variant-overlapping and adjacent fragments. Genes were nominated as candidate targets if a chromatin interaction was observed from a variant-harboring (or adjacent) restriction fragment to the target gene promoter. To map *cis*-regulatory targets of enhancer-promoter interactions, candidate genes initially nominated based solely on physical chromatin interactions with fine-mapped variants were further refined by evaluating whether these variants overlapped melanocytic regulatory regions.

### CRISPRi validation of regulation of RAPGEF1 and UCK1 transcription

To validate regulatory interactions from risk variants to target genes, we performed CRISPR interference (CRISPRi) assays in the immortalized human melanocyte cell line C283T stably expressing dCas9-KRAB [22]. Briefly, C283T cells were infected with a lentiviral vector pLX_311-KRAB-dCas9 (gift from John Doench, William Hahn, and David Root; Addgene plasmid # 96918; http://n2t.net/addgene:96918; RRID: Addgene_96918) followed by monoclonal cell selection. We confirmed stable expression and functional activity of dCas9-KRAB in the clone used for CRISPRi.

Three different guide RNAs (gRNAs) were designed to target the genomic regions flanking each variant site. The gRNA sequences were located ±50bps from the variant (**Supplementary Table 1**). We used two non-targeting gRNAs (NTC1, NTC2) as controls. gRNAs were ligated into the lentiviral vector pXPR-050 (gift from John Doench and David Root, Addgene plasmid #96925; RRID: Addgene_96925). C283T immortalized melanocytes [18] were infected with lentiviral particles encoding gRNA, and at 24h, 1.5 μg/mL of puromycin was added for selection. After two days of puromycin selection, puromycin was removed, and cells were harvested for RNA collection on the same day or one day after puromycin removal. Total RNA was isolated with RNeasy Mini Kit (Qiagen) and cDNA was generated with SuperScript IV VILO Master Mix (Thermo Scientific). Multiple infections were performed with total ten biological replicates for each gRNA against the 3 different variants. mRNA levels of either *RAPGEF1* or *UCK1* were measured by TaqMan assay (Thermo Scientific; **Supplementary Table 1**) and normalized to *GAPDH* levels. qPCR triplicates (technical replicates) were averaged and subsequently considered as a single data point. Average 2^^(delta-Ct)^ values for NTC1 and NTC2 were used for statistical comparisons to other gRNAs, data in tables and graphs are represented as fold-change relative to the average of NTC1 and NTC2. The statistical analysis was performed using a paired two-tailed t-test comparing 2^^(delta-Ct)^ values of individual guides to the average 2^^(delta-Ct)^ of NTC1/NTC2.

### Knock-down of RAPGEF1 and UCK1 expression by shRNA and qRT-PCR quantification of gene expression

Pre-designed shRNAs against human *RAPGEF1* and *UCK1* genes were purchased from Sigma (TRCN0000048129 and TRCN0000048130 for *RAPGEF1*, TRCN0000037763 and TRCN0000199254 for *UCK1*; **Supplementary Table 1**) and expressed in lentiviral vector backbone (pLKO). After viral infection and puromycin selection, total RNA was isolated with the RNeasy Mini Kit (Qiagen) and cDNA was generated with SuperScript IV VILO Master Mix (Thermo Scientific). Transcript levels were measured using TaqMan qPCR assays purchased from Thermo Scientific. *RAPGEF1* and *UCK1* transcript levels were normalized to the levels of *GAPDH*, and PCR triplicates were averaged and considered as one data point.

### Overexpression of RAPGEF1 in human melanocyte culture

*RAPGEF1* was cloned from a cDNA construct (OHS5893-202500127, ORFeome Collab, Horizon Discovery) into pLX304 (a gift from David Root, Addgene plasmid # 25890; http://n2t.net/addgene:25890) or pCW57.1 (a gift from David Root, Addgene plasmid # 41393; http://n2t.net/addgene:41393) lentiviral vectors. This clone is the isoform most strongly expressed across neonatal melanocytes [13]. pCW57.1 contains a doxycycline inducible promoter. Lentiviral vectors were co-transfected into HEK293 cells with the pSPAX2, pMD2-G, and pCAG4-RTR2 packaging vectors. Virus was collected two days after transfection and concentrated by Vivaspin Concentrators (76409-072, VWR). Cells were incubated with virus for 24 h, followed by selection with blasticidin (5-10ug/ml) or puromycin (1–2 μg/ml) for 2–4 days. For pCW57.1-RAPGEF1 infected cells, RAPGEF1 expression was induced by 0.2ug/ml doxycycline in melanocyte culture.

### Gene-based CRISPR-Cas9 knockout screen

A gene-based pooled CRISPR-Cas9 cell proliferation knockout screen was previously performed in an immortalized human melanocyte culture, C283T, as described by Xu and colleagues [14]. Briefly, we designed 3,052 gRNAs covering a set of 288 genes, as well as ⍰200 non-targeting gRNAs. The 288 genes focused primarily on gene candidates from genome-wide significant melanoma GWAS loci. DNA samples were collected from sgRNA infected cells at different times after infection and gRNAs were sequenced and normalized to counts of the integrated gRNA sequence from cells collected prior to Cas9 expression. Prioritized gene hits critical for cell growth and/or survival were called and ranked by using MAGeCK.

### Cell proliferation assays

Cell proliferation was assayed using a BrdU flow kit (BD Pharmingen) according to the manufacturer’s protocol. Briefly, cells were labeled with 10 μM BrdU for 1-3 hours before they were fixed, permeabilized, and subjected to DNase I treatment. Cells were then stained with FITC-conjugated antibody to BrdU and 7-AAD, followed by flow cytometry analysis using an AttuneNxT Cytometer (Thermo Scientific). For crystal violet staining, cells were seeded at equal numbers after infection and drug selection and were stained with crystal violet at different days after seeding. Crystal Violet was solubilized by methanol for quantification as absorbance at 560nm.

### Immunoblotting

For immunoblot analysis, total cell lysates were generated with RIPA buffer (Thermo Scientific) and subjected to water bath sonication. Samples were resolved by 4–12% Bis-Tris ready gel (Invitrogen) electrophoresis. The primary antibodies used were mouse antibody to RAPGEF1 (MA5-26281, ThermoScientific), mouse antibody to RAS (05-516, Sigma), rabbit antibody to RAP1 (ab181858, Abcam), mouse antibody to phospho-ERK(9106S, Cell Signaling Technology), mouse antibody to Pan-AKT (2920s, Cell Signaling Technology), rabbit antibody to total ERK(4695s, Cell Signaling Technology), rabbit antibody to Phospho-AKT(4060s, Cell Signaling Technology), and mouse antibody to β-actin (A5316, Sigma).

### Colony formation assays

Anchorage-independent growth was assayed with p’mel/BRAF^V600E^ cells (provided by H. Widlund, Dana-Farber Cancer Institute). Briefly, 5,000 cells were mixed in HAMF12 medium supplemented with 10% FBS containing 0.4% SeaKem LE agarose (Lonza) and plated on top of a bottom layer with 0.65% agarose prepared in the same HAMF12 medium in a six-well plate. After agarose is solidified, each well was covered with 0.5 ml of feeding medium, which was refreshed twice a week. Colonies were counted and imaged 4-6 weeks after seeding using Lionheart (Agilent) with total magnification of 4x. Colonies were counted from six wells for each condition, and significant differences between samples were assessed by unpaired two-tailed *t*-test assuming equal variance.

### Zebrafish Melanoma Generation and Analysis

Zebrafish melanomas were generated by overexpressing human RAPGEF1 (cloned from a cDNA construct, OHS5893-202500127, ORFeome Collab, Horizon Discovery; same *RAPGEF1* isoform expressed in human melanocyte experiments) or an empty control vector (MCS) in transformed melanocytes using the miniCoopR system [67]. This system involves injection of a Tol2-based miniCoopR vector into the *Tg(mitfa:BRAF*^*V600E*^*); p53(lf); mitfa(lf)* zebrafish strain, where it rescues melanocyte development and simultaneously drives expression of a gene of interest under the *mitfa* promoter. To visualize melanocytes, we co-injected *mitfa:Cas9*, CRISPR gRNAs targeting *tyrosinase* (to block pigmentation), and *mitfa:mCherry*. As melanocyte rescue is mosaic in this system, we screened these cohorts for fish that had *mitfa:mCherry* melanocyte expression in either the RAPGEF1 cohort (n=35), or the MCS control cohort (n=48). These cohorts were comprised of two separate clutches per condition and were genotyped to confirm RAPGEF1 expression status. Starting at 13 weeks post-fertilization, we monitored fish for precursor lesions—termed “cancer precursor zones” (CPZ)—defined as patches of merged *mitfa:mCherry*-high melanocytes exceeding established size and fluorescence thresholds. Screening continued every few weeks until 46 weeks post-fertilization. Kaplan-Meier survival analysis and log-rank (Mantel-Cox) tests showed no significant differences in CPZ onset (p=0.1119) or tumor onset (p=0.211) between groups. Tumor anatomical distribution did not differ substantially between groups.

RAPGEF1 functions upstream of BRAF in the MAPK pathway, and since our primary zebrafish tumor model overexpresses oncogenic *BRAF*^V600E^, this could obscure a role for RAPGEF1 or represent a different biological context to that seen *in vitro*. To assess this, we used the MAZERATI system (Modeling Approach in Zebrafish for Rapid Tumor Initiation) [68] to generate zebrafish cohorts where rescued melanocytes expressed either *BRAFV600E/p53(lf)* or *RAPGEF1/p53(lf)*. In this system, *mitfa:BRAFV600E* and *p53(lf)* mutations are not stably expressed in the zebrafish strain background, but are delivered via plasmid injection at the one-cell stage, allowing us to test whether RAPGEF1 alone can drive tumor initiation in the absence of an activating BRAF mutation. While the *BRAFV600E/p53(lf)* cohort developed tumors at the expected rate, *RAPGEF1/p53(lf)*− fish failed to develop precursor lesions or tumors.

### RAP1 and RAS activation assay

RAP1 activation was measured using the RAP1 activation assay kit from Abcam (ab212011) and RAS activation was measured using GST-fusion Raf-1 RBD beads (Sigma, #14863) or RAF-RBD Protein GST Beads (RF02-A, Cytoskeleton). Cells were grown in supplement-free ATCC medium for 24 hours before treated with 50ng/ml human EGF protein (R&D, Cat# 236-EG-200) at the indicated time. Whole cell extract was taken to be pulled down by the Raf-RBD beads for RAS activation assay or by the RalGDA RBD beads from the Abcam kit for measuring RAP1 activation. Beads were then lysed in SDS loading buffer, resolved by 4–12% Bis-Tris ready gel (Invitrogen) electrophoresis, and blot with RAS (Sigma) or RAP1 (Abcam) antibody. Only the active GTP-bound form of RAS or RAP1 can be pulled down by the beads and the intensity of RAS or RAP1 activation was determined by comparing the pulled-down signal with that from the whole cell extract.

### Assessing RAS-MAPK alterations across multiple melanoma cohorts

We assessed *RAPGEF1* expression association with activating mutations in RAS-MAPK signaling pathway in melanoma tumors. We analyzed cutaneous melanoma tumors across six cohorts: TCGA n=363 [14], Leeds n=503 [69–71], Lund n=146 [72], Liu n=84 [73], Van Allen n=38 [74], Riaz n=39 [75] (**Supplementary Tables 2-8**). Briefly, the Leeds, Lund, and TCGA cohorts included very few samples treated with immunotherapy (TCGA n=23 [76]; Leeds n=12 [70]; and Lund n=4 [72]), while the other three cohorts consisted of patients treated with immunotherapy following tumor collection. Our sample inclusion criteria required melanoma samples that had both mutation and gene expression data available across the cohorts. Cohort details are provided in the **Supplementary Materials**.

### Assessing RAPGEF1 expression association with RAS-pathway

We tested several models classifying RAS-pathway wild-type and mutant tumors, focusing initially on the most commonly-mutated oncogenic driver genes and subsequently adding in less-commonly altered genes. Specifically, within the Leeds sample set (n = 503 total), only specific mutations in *BRAF* (V600|K601) and *NRAS* (Q61|G12|G13) were assessed in a subset of the tumors (n=234), while for the remaining a full set of putative MAPK driver mutations [77] were assessable from sequencing data, so our initial model (Model 1) defined the mutant subgroup as those samples with these specific *BRAF* (V600|K601) and *NRAS* (Q61|G12|G13) hotspot mutations. Additional models were as follows: Model 2 mutant group, any driver mutation in *BRAF* or *NRAS*; Model 3 mutant group, any driver mutation in *BRAF, NRAS, KRAS*, or *HRAS*; Model 4 mutant group, any driver mutation in *BRAF, NRAS, KRAS, HRAS*, or *NF1* (“triple wild-type); and Model 5, any putative driver mutation in the 44 genes defined from the TCGA Pan-Cancer analysis [77] (**Supplementary Table 9**). Outside of the Leeds dataset, all cohorts had full mutation information for *BRAF, NRAS, KRAS, HRAS*, and *NF1*. As described above, Leeds and Lund cohorts had targeted sequencing of select cancer-related genes and were not fully assessed for all genes included in Model 5. In this case for Model 5, we classified wild-type and mutant subgroups based on the available gene sequencing data from each dataset. Methods for annotating individual mutations as drivers are described in the **Supplementary Material**. We assessed an association of *RAPGEF1* levels with RAS-MAPK wild-type and mutant subgroups using a linear regression model, adjusting for age, sex, tumor purity, and tumor stage when available.

### Survival analysis

We conducted survival analyses across multiple melanoma cohorts using the Cox proportional hazards models to assess the association between *RAPGEF1* expression and patient outcomes in tumors wild-type for *BRAF* (V600|K601) or *NRAS* (Q61|G12|G13) hotspot mutations. For each cohort, *RAPGEF1* expression was dichotomized into high- and low-expression groups based on the median expression value of the full dataset. Associations between *RAPGEF1* expression and survival were then evaluated within the wild-type tumor subgroup using the Cox models. To ensure comparability across datasets, all survival times were uniformly converted to days. We performed melanoma-specific survival in the TCGA, Leeds, and Lund cohorts, and overall survival in the Liu, Van Allen, and Riaz cohorts as they lacked melanoma-specific death data. Because stage I tumors were profiled exclusively in the Leeds cohort and are associated with a high survival rate (93-97%) [78], we excluded these tumors from the survival analysis to ensure comparability between the Leeds cohort and other melanoma cohorts, which predominantly included more advanced-stage tumors. For these analyses, we adjusted for covariates specific to each cohort as follows: TCGA: Age, Tumor site; Leeds: Age; Lund: Age; Liu: Cosmic aging; Van Allen: Age. Across all cohorts, the WT tumor subgroup included only a small number of samples, and when the number of events is low relative to the number of covariates, the Cox model can encounter monotone likelihood issues, resulting in extremely large or unstable coefficient estimates. To address this, we applied Firth’s correction.

### Meta-analysis of tumor data

We performed meta-analysis of the *RAPGEF1* expression association with RAS-MAPK subgroups, as well as the association of *RAPGEF1* expression with survival across the analyzed melanoma cohorts as follows: Let be the z-score for cohort. Let ^+^ be the number of subjects with the mutation and ^-^ be the number of subjects of wild type, then the effective sample size 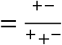. We choose weights 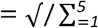 and perform meta-analysis as 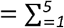. One can show that ~*(0,1)* under _*0*_ and has optimal statistical power assuming equal effect sizes (after normalization).

## Results

### Fine-mapping of the 9q34.13 melanoma risk signal

In order to functionally characterize the melanoma risk locus on chromosome band 9q34.13, which is also associated with nevus count, but not hair color, tanning ability, or telomere length [1], we first assessed whether the locus harbors a single or multiple risk signals. Conditional GWAS analysis using the lead variant at this locus (rs3780269, *P*_GWAS_ =1.92 x 10^-8^, rs3780269-G OR=0.94) revealed no evidence of a secondary signal within +/− 1Mb of the main GWAS association signal (**Supplementary Table 10-11, Supplementary Figure 1A**). Multiple Bayesian fine-mapping methods (SuSiE, DAP-G, and RSparsePro) [25–29] similarly converged on a single and similar credible causal set of variants (CCVs; **Supplementary Table 12**).

To facilitate variant-to-gene mapping, we defined a credible causal set by taking the union of CCVs identified using Bayesian fine-mapping approaches. Within this set, the GWAS lead variant (rs3780269) was consistently assigned the highest posterior probability of being causal across methods (DAP-G PIP = 0.52, RSparsePro PIP = 0.58, SuSiE PIP = 0.55). We also included as CCVs variants identified by fine-mapping directly from GWAS summary statistics by assessing log-likelihood ratio (LLR) relative to the lead variant. Finally, to account for additional variants not genotyped or imputed in the GWAS, we performed LD-based fine-mapping (r^2^ > 0.8 with the lead variant). These methods collectively identified 14 potential causal variants (**Figure 1A, Supplementary Table 12**).

**Figure 1.**
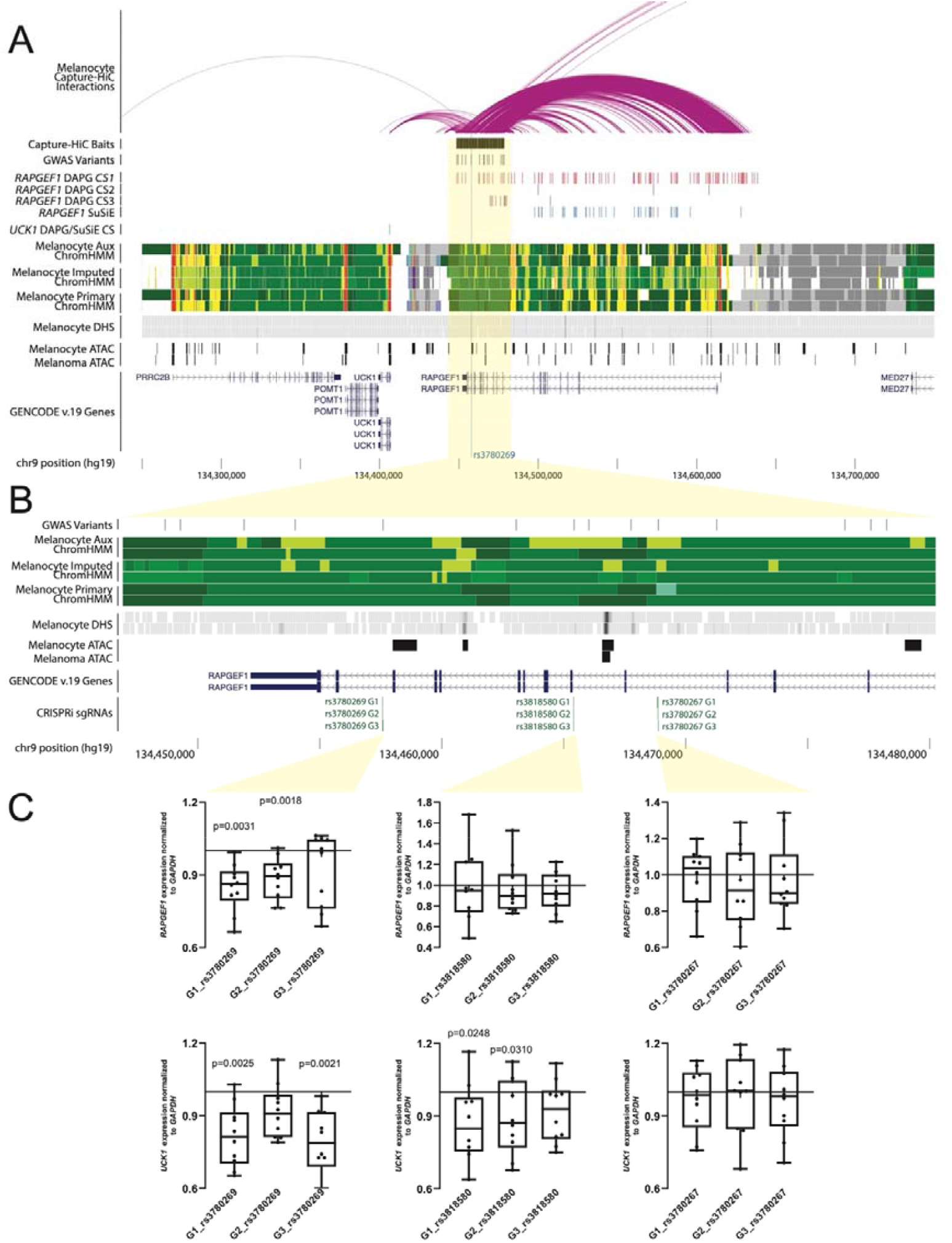
Fine-mapping and epigenomic annotations of melanoma GWAS and melanocyte eQTLs for the 9q34.13 melanoma risk locus. **A**. Genetic variants fine-mapped for the melanoma GWAS are annotated, along with Bayesian fine-mapped variant credible sets (CS) from SuSiE and DAP-G for human primary melanocyte eQTLs for *RAPGEF1* and *UCK1*. Roadmap chromatin states (ChromHMM) from primary melanocytes are shown; red and orange denote annotated promoter, yellow and light green enhancer. Chromatin accessibility from DNAse I hypersensitivity sequencing (DHS) and ATAC-sequencing (ATAC) of melanocytes and melanoma cell models is shown. Chromatin interactions between the entire region of association (Capture-HiC baits) in human melanocytes is shown; fine-mapped variants interact with the promoters of both *RAPGEF1, UCK1*, and *NTNG2* (not shown on figure). **B**. Zoomed in view of the fine-mapped variants. The locations of small guide RNAs used for CRISPR-inhibition (CRISPRi sgRNAs) experiments are shown. **C**. CRISPR-inhibition (CRISPRi) validation of *RAPGEF1 and UCK1* as *cis*-regulatory targets at 9q34.13. Guide RNAs were designed to target three fine-mapped variants within an ~11.3 kb interval. Each guide was individually tested for effects on *RAPGEF1 and UCK1* expression relative to the average of two non-targeting guides in immortalized melanocytes stably expressing dCas9-KRAB via a TaqMan quantitative RT-PCR assay. Whiskers show minimum and maximum values. *P*-values were calculated using a two-sample two-sided paired t-test comparing 2^^(delta-Ct)^ values from individual guides to those from the average of the two non-targeting guides with 10 biological replicates in total.

None of these variants mapped to protein-coding sequence or canonical splice junctions (**Supplementary Table 13**), nor was there a significant or colocalizing splice QTL (sQTL) for rs3780269 (see **Supplementary Material**). One fine-mapped variant, rs3739497 (*P*_*GWAS*_ = 1.9 × 10^−6^; OR = 0.95; risk allele = A; protective allele = G; r^2^ to GWAS lead = 0.88), is located in binding site for two microRNAs (miRNAs; *miR-124-3p* and *miR-506-3p*) in the 3⍰ untranslated region (UTR) of *RAPGEF1*, however we found no evidence that levels of *RAPGEF1* were correlated with either in melanocytes, making it unlikely this variant impacts either miRNA binding or *RAPGEF1* levels (see **Supplementary Material**).

### The 9q34.13 melanoma risk region physically interacts with and regulates RAPGEF1 and UCK1

Several melanoma credible set variants overlap active enhancer or accessible chromatin regions in human melanocytes or melanoma cell models (**Figure 1B, Supplementary Table 14**), while the lead variant is located within a region broadly annotated as an enhancer across multiple cell types (ENCODE *cis*-regulatory element EH38E2732514)[79], suggesting a *cis*-regulatory function. Consistent with this annotation, previous massively parallel reporter assays (MPRA) of fine-mapped risk variants [23] showed the regions surrounding several had significantly different transcriptional activity compared to scrambled DNA sequences in melanocytes and melanoma cells (scramble vs alt comparisons in **Supplementary Table 15**), indicating enhancer and/or repressive function (**Supplementary Figure 2**).

Capture-HiC-based variant-to-gene mapping [22] of the full region of association in primary melanocytes identified chromatin interactions between enhancers harboring fine-mapped variants (**Supplementary Table 14**) and the promoters of *RAPGEF1 and UCK1* (**Figure 1A**; **Supplementary Table 16**), as well as to *NTNG2* (which is lowly-expressed in melanocytes and melanoma). CRISPR-inhibition (CRISPRi) of regions surrounding select variants interacting with *RAPGEF1* and *UCK1* promoters (**Figure 1B**) in immortalized melanocytes confirmed a *cis*-regulatory effect of the region surrounding the GWAS lead variant (rs3780269) for both *RAPGEF1* (2/3 guide RNAs; 0.85–0.88⍰fold relative to non⍰targeting controls, *P* = 0.0031 and *P* = 0.0018 respectively; paired Student’s t-test) and *UCK1* (2/3 gRNAs 0.78–0.82⍰fold, *P* = 0.0025 and *P* = 0.0021 respectively; paired Student’s t-test), as well as rs3818580 for *UCK1* (2/3 guides; *P* = 0.025 and *P* = 0.031) (**Figure 1C**). However, melanocyte and melanoma MPRA data for credible set variants [23] did not find variants with significant allelic differences in transcriptional activity (ref versus alt columns in **Supplementary Table 15**), so while elements in this region regulate both genes, these data do not implicate a specific variant.

### RAPGEF1 and UCK1 are cis-regulatory targets at the 9q34.13 melanoma risk locus in melanocytic cells

Transcriptome wide association studies (TWAS) using eQTL data from human primary melanocytes (n=106) [13] as well as primary melanoma tumors from TCGA (n=103) [14, 15, 80] collectively implicate two genes in the region, with melanoma risk positively associated with predicted melanocyte and melanoma expression of *RAPGEF1* (*P*_melanocyte_ = 4.8 x 10^-5^; *P*_melanoma_ = 4.7 x 10^-5^) and negatively associated with melanocyte *UCK1* (*P*_melanocyte_ = 1.8 x 10^-5^; **Supplementary Table 17**; *UCK1* was not included as a predictable gene from the melanoma dataset). Consistent with TWAS, which considers the collective effects of multiple sequence variants, the risk allele of rs3780269 itself is individually associated with higher *RAPGEF1* in both melanocytes and melanomas (*P* = 1.4 × 10^−11^ and *P* = 5 × 10^−4^, respectively; **Supplementary Figure 3A, Supplementary Table 18, Supplementary Table 19, Supplementary Table 20**) and negatively but not significantly associated with *UCK1* (*P* = 0.22; **Supplementary Figure 3A, Supplementary Table 18**).

In contrast to melanoma GWAS fine-mapping results which suggest a single signal, colocalization and fine-mapping of *RAPGEF1* and *UCK1* eQTL signals point to potentially distinct *cis*-regulatory variants for *RAPGEF1* and *UCK1*, each in partial LD with rs3780269. Melanocyte eQTL results show different lead SNPs for *RAPGEF1* (rs13295485, *P*_*eQTL*_ = 2.8 x 10^-20^, *P*_*GWAS*_ = 4.8 x 10^-5^) and *UCK1* (rs7867616, *P*_*eQTL*_ = 2.8 x 10^-20^; *P*_*GWAS*_ = 3.3 x 10^-5^; **Supplementary Tables 21, 22**), each in partial LD with rs3780269 (1000G EUR r^2^ = 0.50, D’ = 0.74 for rs13295485; r^2^ = 0.17, D’ = 0.66 for rs7867616) and each other (r^2^ = 0.12, D’ = 0.5). Analyses of GWAS and eQTL signals using multiple Bayesian approaches (SuSiE + COLOC [54] and SharePro [55]) do not provide support for colocalization of these eQTLs with the melanoma GWAS (**Supplementary Tables 23, 24; Supplementary Figure 3B, C**). Fine-mapping the *RAPGEF1* eQTL in melanocytes reveals multiple sets of CCVs collectively containing over 100 variants and spanning the entire gene and promoter (**Figure 1A, Supplementary Table 25**); critically several overlap the set of melanoma CCVs (**Supplementary Table 12**). The single fine-mapped *UCK1* eQTL credible set variant overlaps annotated *UCK1* promoter (rs7867616, **Figure 1A, Supplementary Table 26**) but not the melanoma credible set. Consistent with the eQTL data, we note a strikingly similar pattern in melanocyte methylation QTL (meQTL) data (see **Supplementary Material**), where a methylome-wide association study identified significant CpG methylation probes in both genes and individual meQTL signals mirror eQTL signals for the respective genes. These data collectively reflect a complex *cis*-regulatory structure underlying the 9q34.13 association signal.

### Contributions of RAPGEF1 and UCK1 expression to the melanoma association signal

To further reinforce *RAPGEF1* and or *UCK1* as strong causal candidates regulated by a complex pattern of *cis*-regulatory signals, we performed additional conditional GWAS analyses. Conditioning on predicted melanocyte *RAPGEF1* and *UCK1* expression each partially attenuated the GWAS signal for lead SNP rs3780269 (*P*_GWAS_ = 1.9 x 10^-8^, *RAPGEF1 P*_conditional_ = 1.0 x 10^-4^; *UCK1 P*_conditional_ = 2.1 x 10^-5^; **Supplementary Table 27, 28**). When both genes were included in a joint model, both are significant (**Supplementary Table 29**) and the signal for rs3780269 is dramatically attenuated (*P*_conditional_ = 0.002; **Supplementary Table 30**). We observe similar results when instead conditioning on respective lead eQTL SNPs (*RAPGEF1 P*_conditional_ = 4.9 x 10^-3^, *UCK1 P*_conditional_ = 6.33 x 10^-5^; **Supplementary Table 31, 32; Supplementary Figure 1B, 1C)**, suggesting a contribution of *cis*-regulatory variants for both genes to the melanoma risk signal. We note that conditioning only on the fine-mapped *RAPGEF1* eQTL SNP with the highest PIP identified by DAP-G itself resulted in complete attenuation of the GWAS signal (*P*_conditional_ = 0.29, **Supplementary Table 33, Supplementary Figure 1D**), perhaps suggesting a critical role for *RAPGEF1*. Taken together, these results point to multiple causal variants in partial LD to rs3780269 respectively influencing two plausible effector genes, *RAPGEF1* and *UCK1*, with both genes having opposing effects correlated with melanoma susceptibility: decreased *UCK1* expression and increased *RAPGEF1* expression are linked to risk.

### RAPGEF1 and UCK1 both are critical for melanocyte growth and overexpression of RAPGEF1 stimulates melanocyte growth

Given the opposing effects for *RAPGEF1* and *UCK1* on melanoma risk, we assessed whether their expression impacts melanocyte growth and/or survival. We previously included *RAPGEF1*, but not *UCK1*, in a small pooled CRISPR proliferation screen in human melanocytes, noting the set of guides targeting *RAPGEF1* as the third most depleted amongst all genes screened (*P* = 4.97 x 10^-6^) [18] suggesting a strong positive role in melanocyte proliferation and or viability. Here, we validated this finding, silencing *RAPGEF1* using two independent shRNAs in immortalized human melanocytes and assessing cell proliferation by crystal violet staining and BrdU incorporation assays. Knockdown of *RAPGEF1* using both guides reduced melanocyte number as assessed by crystal violet assay (**Figure 2A, 2B**). Consistent with these data, BrdU flow cytometry revealed a significant decrease in DNA synthesis after targeting with both shRNAs, indicating a block in cell cycle progression upon *RAPGEF1* knockdown (**Figure 2C**). Parallel knockdown of *UCK1* similarly resulted in reduced proliferation in immortalized melanocytes, as evidenced by both assays (**Figures 2D-F**), mirroring the effects observed with *RAPGEF1* suppression. Critically, while a positive role for *RAPGEF1* in melanocyte proliferation is consistent with observation that increased melanocyte *RAPGEF1* is associated with melanoma risk, the essential role of *UCK1* in melanocyte proliferation contrasts with eQTL and TWAS findings which show that increased *UCK1* expression is protective against melanoma. While we cannot exclude a potential role for *UCK1* in melanoma susceptibility at the 9q34.13 locus via means other than influencing melanocyte proliferation, the strong concordance between the essential function of *RAPGEF1* in melanocyte proliferation and its elevated expression linked to increased melanoma risk led us to prioritize *RAPGEF1* for further investigation.

**Figure 2.**
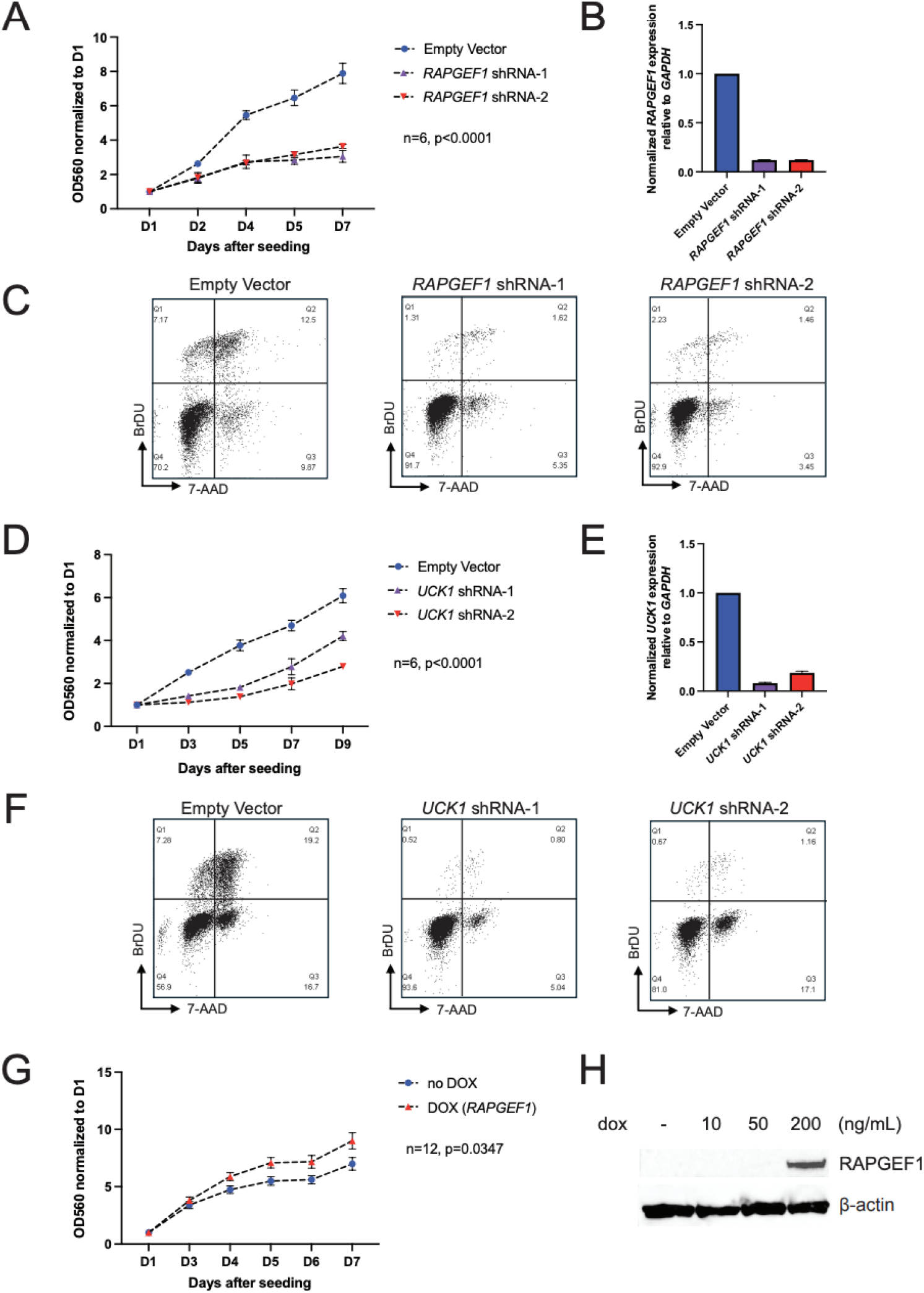
RAPGEF1 and UCK1 positively regulate melanocyte proliferation. **A**. Crystal violet assays following *RAPGEF1* knockdown in immortalized melanocytes. Equal numbers of cells either infected with two *RAPGEF1* shRNAs or vector control were seeded after infection and drug selection. Cells were fixed and stained with crystal violet at multiple timepoints after seeding. Crystal violet was then solubilized from stained cells and measured by OD560. OD560 values were normalized to the reading at D1, and the mean of six biological replicates and SEM were plotted. *P*-value was calculated from two-way ANOVA; **B**. Quantitative RT-PCR for *RAPGEF1* in immortalized melanocytes following infection with shRNAs targeting *RAPGEF1* relative to empty vector control. RNA was collected and assayed at D5 after infection. **C**. FACS analysis of immortalized melanocytes infected with either empty vector or *RAPGEF1* shRNA at D5 after infection. BrdU and 7-AAD staining were used to characterize the cell cycle status of different cells. The images shown here are from a representative experiment from three biological replicates. **D**. Crystal violet assays following *UCK1* knockdown in immortalized melanocytes. Equal numbers of cells either infected with two *UCK1* shRNAs or vector control were seeded after infection and drug selection. Cells were fixed and stained with crystal violet at multiple timepoints after seeding. Crystal violet was then solubilized from stained cells and measured by OD560. OD560 values were normalized to the reading at D1, and the mean of six biological replicates and SEM were plotted. *P*-value was calculated from two-way ANOVA. **E**. Quantitative RT-PCR for *UCK1* in immortalized melanocytes following infection with shRNAs targeting *UCK1* relative to empty vector control. RNA was collected and assayed at D7 after infection **F**. FACS analysis of immortalized melanocytes infected with either empty vector or *UCK1* shRNA at D7 after infection. BrdU and 7-AAD staining were used to characterize the cell cycle status of different cells. The images shown here are from a representative experiment from three biological replicates. **G**. Immortalized melanocytes infected with a dox-inducible vector (pCW57.1) to express *RAPGEF1* were seeded after infection and drug selection. Dox-treated cells were treated with 200ng/mL dox at seeding (D0) and cells were fixed and stained with crystal violet at multiple timepoints after seeding. Crystal violet was then solubilized from stained cells and measured by OD560. OD560 values were normalized to the reading of D1 and the mean of 12 biological replicates and SEM were plotted. *P*-value was calculated using two-way ANOVA. **H**. Western blot for RAPGEF1 in dox-treated immortalized melanocytes. Whole cell extract was collected 48 hours post-dox treatment at D7 after infection.

We then assessed whether increased *RAPGEF1* levels contributed to increased proliferation in immortalized melanocytes using a doxycycline induced promoter construct. Upon doxycycline treatment, cells showed significantly enhanced growth by crystal violet staining compared to untreated controls (**Figure 2G, H**), demonstrating that increased *RAPGEF1* expression stimulates melanocyte growth and could contribute to melanoma development. In summary, *RAPGEF1* expression influences melanocyte proliferation, with elevated levels driving increased cell growth that could in turn contribute to nevus and or melanoma development.

### RAPGEF1 overexpression triggers malignant transformation of human melanocytes

Having established that *RAPGEF1* overexpression enhances melanocyte growth, we next investigated whether *RAPGEF1* functions as a potential oncogene capable of promoting malignant transformation. While RAPGEF1 has diverse roles in cancer depending on the cellular context, tumor type, and stage, its function as a GEF for the small GTPase protein RAP1 is central to its potential oncogenic activity. Mechanistically, RAP1 can initiate and maintain MAPK/PI3K signaling, which can trigger malignant transformation of cells. To further evaluate whether *RAPGEF1* can stimulate malignant transformation by affecting melanocyte growth, we examined the potential effect of RAPGEF1 on anchorage-independent growth of TERT-immortalized human melanocytes (p’mel cells) by soft-agar colony formation assay. While p’mel cells themselves do not efficiently form large colonies in soft-agar, stably expressing BRAF^V600E^ in p’mel together with other oncogenes lead to colony formation [81, 82]. NRAS and its constitutively active mutants have been shown to promote malignant cellular transformation in multiple cells of different origins [83, 84] while the direct role of RAP1 in cellular transformation has not been well-reported. Here, we observed enhanced colony formation (>70 µm) in p’mel cells stably expressing both BRAF^V600E^ and constitutively activated NRAS, specifically NRAS^G12V^ or NRAS^Q61R^ (*P* < 0.0001, P < 0.0001, respectively; **Figure 3A, B**) relative to those expressing BRAF^V600E^ alone. Not surprisingly, enhanced colony formation was also observed in p’mel cells stably expressing both BRAF^V600E^ and a synthetic constitutively activated RAP1A^G12V^ mutant (P < 0.0001). By introducing RAPGEF1 overexpression in p’mel cells, we observed that RAPGEF1 similarly cooperated with *BRAF*^*V600E*^ to enhance colony formation (*P* < 0.0001). The NRAS^Q61R^ mutant induced a stronger colony formation effect than the NRAS^G12V^ mutant (*P* < 0.0001), consistent with *NRAS* mutations at codon 12 (such as G12V) generally exhibiting intermediate oncogenic potency compared to the stronger activating mutations at codon 61 [85]. The number of colonies formed upon *RAPGEF1* overexpression was lower than those induced by NRAS^Q61R^ (*P* < 0.0001) and RAP1A^G12V^ (*P* = 0.0002), but on a more similar level to wild-type RAP1A (*P* = 0.10) or NRAS^G12V^ (*P* = 0.32; **Figure 3A, B**). We note that we did not observe such cooperation in a BRAF^V600E^-expressing zebrafish model (*Tg(mitfa:BRAF(V600E));⍰p53*^−*/*−^) [67], where overexpression of human RAPGEF1 did not accelerate formation of cancer precursor zones (CPZs; *P* = 0.11) or tumors (*P* = 0.21), nor was RAPGEF1 overexpression sufficient to drive tumor formation in fish that did not overexpress BRAF^V600E^. Overall, these data suggest that *RAPGEF1* possesses an intermediate level of oncogenic potential, contributing to colony formation in melanocytes but insufficient to drive tumorigenesis in the absence of a strong oncogenic mutation *in vivo*.

**Figure 3.**
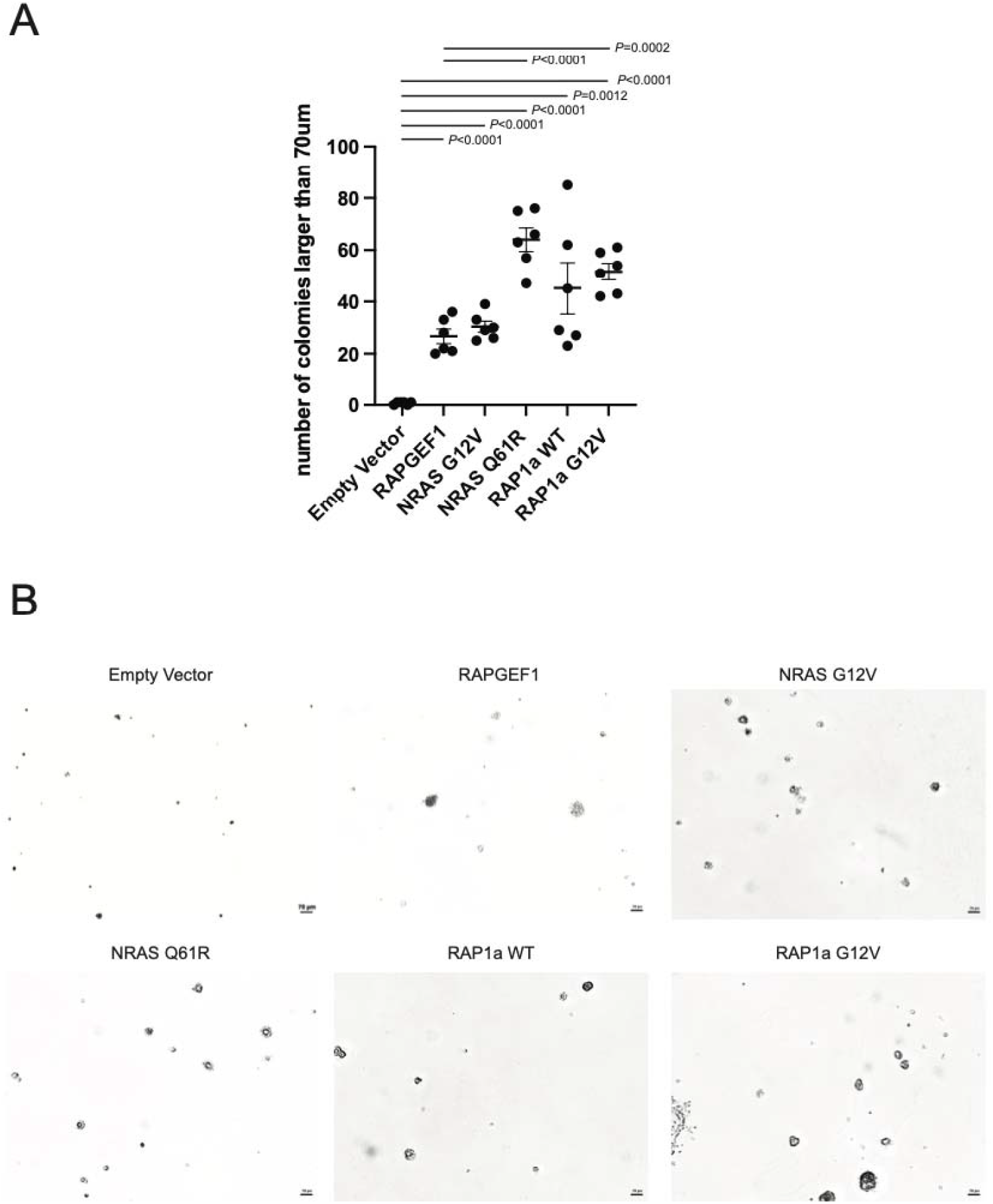
Colony formation in soft agar of p’mel cells stably expressing *BRAF*^*V600E*^ infected with lentivirus to express either RAPGEF1, NRAS^G12V^, NRAS^Q61R^, RAP1A, or RAP1A^G12V^. **A**. Immortalized melanocytes (p’mel cells) expressing constitutively active *BRAF*^V600E^ were infected with empty pLX304 vector or constructs expressing different proteins and seeded in soft-agar plates after blasticidin selection. Images were taken between week 4-5 after plating, and colonies were counted from two sets of triplicates (*n* = 6), with counts from each individual replicate shown as a dot. Whiskers show mean with SEM. *P*-values were calculated using two-tailed, unpaired Student’s t-test. Significance is shown only for comparisons with the empty vector control and RAPGEF1 overexpression. **B**. Representative images of colonies are shown. The same magnification (4x) was applied for all the images.

### RAPGEF1 overexpression in serum-starved melanocytes potentiates activation of both RAS and RAP1

Since melanocyte growth and malignant transformation effects were noted for RAPGEF1 overexpression, we hypothesized that its overexpression promotes oncogenic activity via activation of its known target RAP1. To test this, we overexpressed RAPGEF1 in melanocytic cells using a tetracycline-inducible system, and RAP1 activation was assessed by pulldown with RalGDS RBD beads, which selectively bind to the GTP-bound active form of RAP1. Under standard melanocyte culture conditions, RAP1 activation remained minimal regardless of RAPGEF1 overexpression, suggesting that cell growth supplement factors may suppress or mask RAPGEF1 related effects (data not shown). However, when cells were grown in supplement-free medium for 24 hours and then stimulated with human Epidermal Growth Factor (hEGF), a weak, transient increase in RAP1 activation was detected in cells not treated with doxycycline (dox), while dox-treated RAPGEF1 overexpressing cells showed enhanced activation, with a marked increase in active RAP1 at 15 minutes following hEGF treatment (**Figure 4A**). These results are consistent with previous studies showing that RAPGEF1 acts as a GEF for RAP1 in mammalian cells.

**Figure 4:**
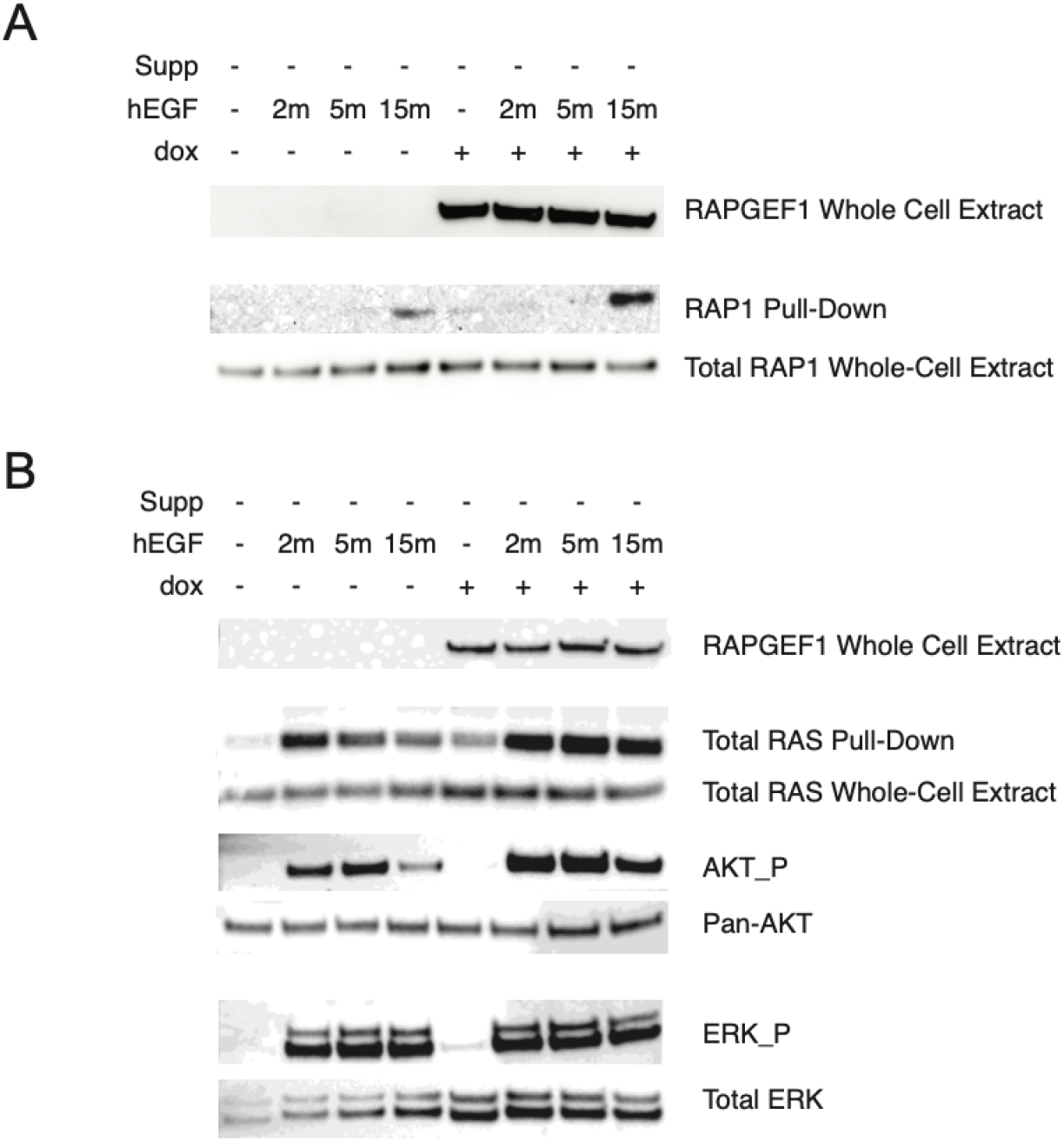
RAPGEF1 potentiates RAP1 and RAS activation in serum starved melanocytes. C283T immortalized melanocytes with doxycycline-inducible expression of *RAPGEF1* were grown in supplement-free medium for 24hrs and subsequently treated with human EGF (hEGF) protein for the indicated time. **A**. Active GTP-bound RAP1 was pulled down by RalGDS-RBD beads from whole-cell extract. Pull-down and whole-cell (total RAP) fractions were compared by Western blotting with an antibody against RAP1. **B**. Active GTP-bound RAS was pulled down by Raf-RBD beads, with the pull-down and whole-cell (total RAS) fractions compared by Western blotting with an antibody against RAS. Active and total AKT and ERK were compared via Western blotting. Figures are representative experiments from three biological replicates.

RAP1 belongs to RAS small G-protein family which also contains NRAS, HRAS, KRAS and many other RAS proteins. Almost all known GEFs for RAS family G-proteins contain a CDC25 homology domain as the catalytic domain, yet each GEF acts on only a small subset of the RAS family members with a high selectivity [86]. While RAPGEF1 has been established as a major GEF for RAP1, its role as a GEF of other RAS family proteins is controversial. *In vitro* GEF assays using RAPGEF1 containing only CDC25 domain showed no activation of any RAS proteins [86] whereas full-length RAPGEF1 displayed activation of R-RAS both *in vitro* and in living COS7 cells [87]. At the same time, it was reported that RAP1 activation by RAPGEF1 might also contribute indirectly to RAS activation [88]. Given that the RAS signaling pathway is a key regulator of melanoma development, RAS alterations are common in melanoma, and that nearly every nevus harbors a mutation in BRAF or RAS, we asked whether RAPGEF1 overexpression affects RAS activation in melanocytes as we observed for RAP1. Compared to controls, RAPGEF1 overexpression led to RAS activation even in the absence of hEGF, with stronger RAS activation observed in RAPGEF1 overexpressing cells following hEGF stimulation (**Figure 4B**). We also assessed downstream signaling activity following RAS activation, noting increases in ERK and AKT activation correlating with levels of activated RAS (**Figure 4B**). Notably, although RAPGEF1 overexpression enhanced both RAP1 and RAS activation in response to hEGF, the kinetics of activation between RAS and RAP1 is clearly different, with strong RAS activation from 2 to 15 minutes in comparison to noticeable active RAP1 only at 15 minutes post hEGF treatment. These data raise the possibility that the activation of RAS and RAP1 by RAPGEF1 may not happen concurrently, and may be subjected to spatio-temporal regulation [89]. These data support a model in which dysregulated RAPGEF1 expression amplifies not only RAP1, but also RAS signaling in melanocytes, driving abnormal melanocyte proliferation and survival and contributing to melanogenesis and tumor development.

### Elevated RAPGEF1 levels are associated with tumors lacking activating mutations in RAS-MAPK signaling

Hyperactivation of the RAS-MAPK signaling is a hallmark of melanoma [90, 91]. Most melanomas acquire activating mutations in *BRAF* or *NRAS*, or loss-of-function mutations in *NF1*, which result in constitutive RAS-MAPK pathway activation [92, 93]. In tumors lacking these canonical drivers, rare mutations in other RAS-MAPK pathway members have been reported to sustain RAS-MAPK signaling [14, 94–96]. In melanocytes, RAPGEF1 overexpression induced RAS activation in the absence of hEGF treatment. Based on this, we hypothesized that *RAPGEF1* expression may act as an alternative contributor to driving RAS-MAPK activation in melanoma tumors without known or strongly-activating RAS-MAPK activating alterations. To test this, we analyzed RAPGEF1 expression in six publicly available melanoma cohorts that included both mutation and gene expression profiling data (TCGA melanoma n=366 [14], Leeds n=503 [69–71], Lund n=146 [72], Liu n=84 [73], Van Allen n=38 [74], Riaz n=39 [75]; **Supplementary Tables 2-8**). Since mutational data for specific strongly-activating *BRAF* (codons V600 and K601) and *NRAS* (codons Q61, G12, and G13) alterations across the widest range of tumors across all datasets (a subset of the Leeds cohort was only assessed for these specific mutations), we initially focused on defining the RAS-MAPK wild-type and mutant groups based on these specific alterations. After adjusting for clinical and technical covariates (age, sex, tumor stage, and tumor purity), we consistently observed significantly elevated *RAPGEF1* expression in wild-type tumors compared to the mutant subgroup across five of six cohorts (TCGA *P* = 3.4 x 10^-4^; Leeds *P* = 0.01; Lund *P* = 0.007; Liu *P* = 0.004; Van Allen *P* = 0.02), with the direction of effect in the non-significant Riaz cohort consistent with the other five (*P* = 0.2) (**Supplementary Table 34; Figure 5A**). Meta-analysis across these datasets showed a highly significant difference (P_meta_ = 6.6 x 10^-10^). We also expanded to consider models cumulatively adding additional *BRAF* and *NRAS* driver mutations (Model 2), *HRAS* and *KRAS* driver mutations (Model 3), loss-of-function *NF1* mutations (Model 4), and driver alterations in other RAS-MAPK pathway members using definitions from the TCGA Pan-Cancer analysis [77] (Model 5; models listed in **Supplementary Table 9**). *RAPGEF1* expression remained consistently higher in the wild-type subgroups in meta-analyses of each model, with the strongest association in models considering driver mutations specifically in *BRAF* and *NRAS* (Model 2; *P*_model2_ = 4.54 x 10^-10^; *P*_model3_ = 1.09 x 10^-8^, *P*_model4_ = 2.73 x 10^-6^, *P*_model5_ = 1.96 x 10^-3^; **Supplementary Table 34**). Taken together, the consistent enrichment of *RAPGEF1* expression in tumors lacking RAS-MAPK activating alterations, coupled with its ability to induce RAS, RAP, and MAPK activation in melanocytes, supports a model in which *RAPGEF1* contributes to RAS-MAPK pathway activation in the absence of strongly-activating canonical driver mutations.

**Figure 5:**
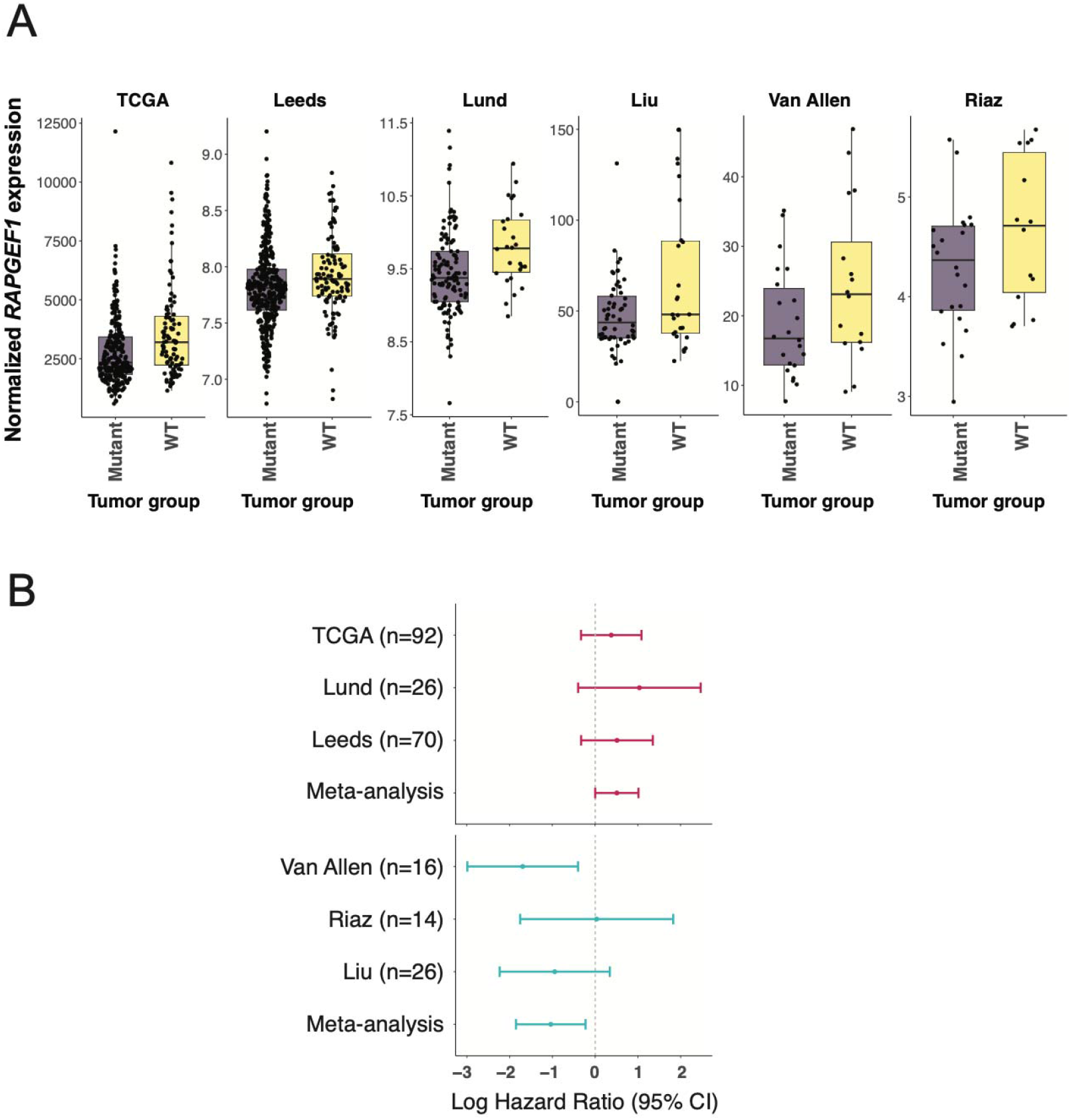
*RAPGEF1* is consistently higher in NRAS/BRAF-wild type tumors across six independent melanoma tumor cohorts. **A**. We assessed mutation and RNA expression data from six melanoma tumor cohorts: TCGA, Leeds, Lund, Liu, Riaz, and Van Allen cohorts. Within each cohort, melanoma tumors were classified into two groups: mutant and wild-type (WT) based on the presence of any *BRAF* V600 or K601 or *NRAS* Q61, G12, or G13 codon somatic mutations; tumors were designated wild-type otherwise. For each cohort, *RAPGEF1* expression distribution is visualized using boxplots. In each boxplot, the central black line represents the median expression value, upper whisker extends from the third quartile (Q3) to the largest value within 1.5x inter-quartile range (IQR) above Q3, while the lower whisker spans from the first quartile (Q1) to the smallest value within 1.5⍰×⍰IQR below Q1. The interquartile range (IQR) is defined as the difference between Q3 and Q1. Because expression data were normalized using a cohort-specific normalization method, *RAPGEF1* expression values are not directly comparable between cohorts. **B**. Meta-analysis across melanoma cohorts assessing *RAPGEF1* as a marker of survival in tumors wild type for *BRAF* codon V600 and K601 and *NRAS* codon Q61, G12, and G13 somatic mutations. To minimize potential confounding due to treatment-specific effects, we analyzed datasets comprised primarily of patients not treated with immunotherapy (TCGA, Lund, and Leeds cohorts) separately from those focused on immunotherapy-treatment (Van Allen, Riaz, and Liu cohorts). The variable n denotes the number of samples analyzed in the WT group for each cohort. Within each cohort, *RAPGEF1* expression was dichotomized into high and low groups based on the median value. The low *RAPGEF1* expression group is the reference group in Cox models and is depicted by the grey line on the plot. Each point represents the estimated log HR for a given cohort; horizontal lines denote the 95% confidence interval around the point estimate. The meta-analysis label on the plot denotes the log HR estimate from the meta-analysis.

### RAPGEF1 expression is associated with prognosis in tumors lacking RAS-MAPK activating alterations

We next evaluated whether *RAPGEF1* expression has prognostic significance in melanoma tumors lacking RAS-MAPK activating alterations. Since *BRAF* (V600|K601) and *NRAS* (Q61|G12|G13) hotspot alterations were profiled across most cohorts; we limited survival analysis to tumors that were wild-type for these alterations. To minimize potential confounding due to treatment-specific effects, we analyzed datasets comprised primarily of patients not treated with immunotherapy separately from those focused on immunotherapy-treatment. The Leeds, Lund, and TCGA cohorts predominantly included patients who had not received immunotherapy, whereas the Liu, Van Allen, and Riaz cohorts comprised immunotherapy-treated patients. *RAPGEF1* expression was dichotomized into high and low groups based on a cohort-wide median cut-off for each cohort. In the analysis, non-immunotherapy-treated WT tumors from TCGA (n = 92), Leeds (n = 70), and Lund (n = 26) cohorts, higher *RAPGEF1* expression subgroup was associated with poorer prognosis compared to *RAPGEF1* low expression group (TCGA HR = 1.5, Leeds HR = 1.7, Lund HR = 2.8; **Supplementary Table 35, Figure 5B**). Though these associations are below significance threshold within individual cohorts, a meta-analysis of these cohorts is significant (P=0.049) for *RAPGEF1* overexpression predicting shorter survival (HR = 1.6; **Supplementary Table 35; Figure 5B**). In contrast, for cohorts of patients with WT tumors treated with immunotherapy (Liu = 26, Van Allen = 16, Riaz = 14), the *RAPGEF1* high expression group was associated with improved prognosis (Liu HR = 0.4, Van Allen HR = 0.2, Riaz HR = 1.03) and was significant upon meta-analysis of these cohorts confirming a significant association between higher *RAPGEF1* expression and longer survival (HR = 0.4, *P* = 0.01; **Supplementary Table 35, Figure 5B**). The opposing effects of *RAPGEF1* expression association between relatively non-treated and treatment cohorts is noteworthy, however given the low sample size across most cohorts, these preliminary findings should be validated in larger, independent cohorts to confirm and extend these observations.

## Discussion

GWAS of melanoma and melanoma risk-associated phenotypes have successfully identified numerous loci associated with risk, but the causal sequence variants and genes underlying most have yet to be established. Here, we characterize a GWAS locus at 9q34.13 associated with melanoma risk as well as nevus count [1] and identify a likely role for germline variation regulating RAS and RAP signaling via modulation of expression of the guanine nucleotide exchange factor *RAPGEF1*. Given that both nevi and melanomas originate from melanocytes, we focused on quantitative trait locus (QTL) data from melanocytic cells, demonstrating an association between higher *RAPGEF1* expression and the risk allele via both colocalization and TWAS/MWAS approaches. We show that RAPGEF1 promotes both RAP1 and RAS activation in melanocytes and contributes to colony formation in immortalized melanocytes. Notably, we also observe consistently higher *RAPGEF1* levels in melanomas lacking canonical, strongly-activating RAS-MAPK pathway mutations, suggesting that *RAPGEF1* could potentially contribute to pathway activation in these wild-type tumors.

Analyses of this locus using multiple melanocytic genomic datasets (eQTL, meQTL) revealed a complex pattern of LD between the melanoma GWAS signal and two partially-correlated melanocyte eQTLs in the region for *RAPGEF1* and *UCK1*, complicating efforts to firmly identify the causal sequence variant or variants underlying melanoma risk at this locus. While conditional analysis of this GWAS locus indicates the presence of only a single signal, fine-mapping of the GWAS and respective melanocyte eQTLs reveal mostly distinct sets of potential causal variants for each, all in LD with each other, possibly indicating the presence of functional variants in each credible set contributing to risk, or alternatively that the GWAS signal may simply be an efficient tag for haplotypes containing causal *UCK1* and *RAPGEF1 cis*-regulatory variants. We demonstrated that the region harboring the GWAS signal itself physically interacts with the promoters of and is a functional regulatory sequence for both genes. Conditioning on predicted melanocyte expression of both *RAPGEF1* and *UCK1* together does not fully attenuate the GWAS signal, suggesting a potential functional role for the GWAS signal itself. Still, past MPRA work assessing fine-mapped melanoma GWAS variants in both melanocytes and melanoma failed to identify variants with allele-specific *cis*-regulatory activity, perhaps indicating the lack of causal variants amongst the GWAS credible set or alternatively that there may be undiscovered variation strongly associated with the signal, or both. Further work will be required to identify the causal *cis*-regulatory variants in this region.

Similarly, the complex LD between GWAS and eQTL signals in this region complicate efforts to firmly establish a single gene as the only likely causal gene underlying melanoma risk at this locus. eQTL and meQTL analyses establish both *RAPGEF1* and *UCK1* as *cis*-regulatory targets and strong candidate melanoma risk genes where higher *RAPGEF1* expression and lower *UCK1* are associated with risk. Conditional analysis of the GWAS based either on predicted *RAPGEF1* and *UCK1* expression, or alternatively using lead eQTL SNPs for both genes showed that *RAPGEF1* expression potentially contributes more to the melanoma risk signal than *UCK1*, but these data need to be interpreted cautiously given potential differences in the populations from the GWAS and the relatively small sample of melanocytes and melanomas used in QTL analyses.

We performed gene knockdown experiments in melanocytes to assess potential phenotypes mediated by both genes. Knockdown of either gene resulted in dramatically decreased melanocyte proliferation. Given the opposing directions of effect for melanocyte eQTLs relative to melanoma risk, we chose to focus on *RAPGEF1* in melanocytes where higher expression associated with risk and an observed positive effect on melanocyte growth is consistent with a role promoting tumor formation or growth. Nonetheless, we cannot fully exclude a role for *UCK1*, a member of the pyrimidine salvage pathway that is crucial for DNA replication and repair mechanisms. We observed that RAPGEF1 overexpression in an immortalized melanocyte model that expresses oncogenic BRAF (V600E) results in efficient formation of large (>70um) colonies, where BRAF^V600E^ expression itself is not sufficient. These data are consistent with that observed for other genes with oncogenic potential in melanocytes in this model [81, 82]. While colony formation in melanocytes expressing constitutively active mutants of NRAS (Q61R) and RAP1A (G12V) was considerably higher than that for RAPGEF1, colony formation was similar between RAPGEF1 overexpressing melanocytes and those expressing the less common and potent G12V mutant of NRAS, as well as overexpression of wild-type RAP1A, highlighting the oncogenic activity of RAPGEF1.

*RAPGEF1* is a guanine nucleotide exchange factor that facilitates the exchange of GDP for GTP, thereby activating small GTPases, through interactions mediated by SH3 domains of its partner proteins. Almost all known GEFs for RAS family G-proteins contain a CDC25 homology domain as the catalytic domain, yet each GEF acts on only a small subset of the RAS family members with a high selectivity [86]. While SOS proteins are widely-expressed GEFs activating RAS proteins, RAPGEF1 has been established as a major GEF for RAP1 [97, 98]. Given the similarity in RAPGEF1 colony formation results as compared to NRAS^G12V^, and that RAPGEF1 has previously been implicated in activation of R-RAS [87] as well as indirect activation of RAS [88], we assessed the effect of RAPGEF1 expression on both RAP1 and RAS activation in serum starved melanocytes. Strikingly, RAPGEF1 overexpression contributed to both RAP1 and RAS activation upon hEGF treatment, as well as leading to constitutive activation of RAS itself in the absence of hEGF suggesting RAPGEF1 contributes to RAS activation. Temporally, we observe that RAS activation in response to hEGF appears to precede activation of RAP1, with a noticeable increase in active RAS at 2 minutes compared to that for RAP1 at 15 minutes.

While a role for direct activation of RAS by RAPGEF1 has not been established, some GEFs can indeed act on both RAS and RAP1 proteins. GRP3, a GEF abundantly expressed in brain, increased GTP/GDP-bound ratios of both HRAS and RAP1A in 293T cells [99]. CalDAG-GEFIII, one of the four CalDAG-GEFs, promoted nuclear exchange of HRAS, R-RAS and RAP1, and shows the widest substrate specificity among the known GEFs [100]. RasGRP2, an alternatively spiced form of CalDAG-GEF1, selectively catalyzes nucleotide exchange on N- and K-RAS, but not H-RAS. RasGRP2 also activates RAP1, albeit less potently than CalDAG-GEF1, which preferentially activates RAP1 [101]. The selectivity of a substrate by a GEF is largely determined by the amino acid residues at the interface between the GEF and the GTPase, yet this interaction can be highly context-dependent and subjected to spatio-temporal regulation [89, 102]. During *Drosophila* eye and wing development, overexpression of membrane-tagged full-length *Drosophila* RAPGEF1 phenotypically mimics overactivation of the RAS1–MAPK pathway, suggesting that *Drosophila* RAPGEF1 is involved in RAS activation *in vivo*. These effects can be suppressed by both reduction in the gene dose of RAS1–MAPK pathway components or RAP1, as well as overexpression of a membrane-tagged version of *Drosophila* RAPGEF1 lacking the CDC25 catalytic domain. These findings indicated that membrane localization of dysregulated RAPGEF1 can trigger activation of RAP1 and RAS resulting in the activation of MAPK [102]. We did perform mass spectrometry analysis of proteins pulled-down by RAPGEF1 antibody in RAPGEF1 overexpressing immortalized melanocytes grown either in regular melanocyte medium or under serum-starved and hEGF-treated conditions, but did not identify any small GTPases (including RAP1; data not shown). While the traditional affinity-purification Mass spectroscopy (AP-MS) methods often require stable protein interactions in cell extracts, small GTPases undergo dynamic conformational changes as they cycle between active and inactive states and the transient nature of their interactions with GEFs making it difficult to be captured. A typical example is SOS1, a major GEF of RAS protein, is historically challenging to be captured with AP-MS [103, 104]. Interestingly, using live-cell proximity-dependent biotin labelling of proteins (BioID) followed by Mass Spec analysis, Kovalski and colleagues captured an interaction between HRAS and RAPGEF1 in bladder cancer cells, supporting the notion that RAPGEF1 may indeed function as a GEF of HRAS [104]. Further work will be required to assess whether RAPGEF1 directly acts on RAS in melanocytes, however our data clearly demonstrate a role for activation of RAS by RAPGEF1 in melanocytes whether directly or indirectly.

Most melanoma tumors harbor somatic RAS-MAPK pathway activating alterations, including oncogenic mutations in RAS genes, BRAF, or loss of NF1, and likely do not need to rely on more subtle germline influence on pathway-activating genes. Still, a minority of melanomas lack strongly-activating RAS and BRAF mutations or alterations in other directly-involved pathway members, suggesting other mechanisms likely contribute to pathway activation in a subset of tumors. Given our observation that RAPGEF1 activates RAS in melanocytic cells, we evaluated whether *RAPGEF1* expression differs between tumors with RAS-MAPK activating mutations and those that are wild-type. Across six melanoma cohorts, we consistently observed elevated *RAPGEF1* expression in wild-type tumors. These data are consistent with a potential role for RAPGEF1 contributing to RAS-MAPK pathway activation in these tumors, whether via an effect on RAS, RAP, or both. We also assessed whether *RAPGEF1* expression in wild-type tumors could serve as a prognostic marker. Across melanoma cohorts where most patients had not received any form of immunotherapy treatment, *RAPGEF1* overexpression predicted poor prognosis. Conversely, in cohorts where all patients were treated with immunotherapy, elevated *RAPGEF1* expression predicted improved survival. Interestingly, rare somatic mutations in *RAPGEF1* have been observed in non-Hodgkin’s lymphomas that disrupt RAPGEF1 autoinhibition and lead to constitutive RAP1 activation, suggesting a plausible role for RAPGEF1 in promoting cancer progression [105]. Nonetheless, given the relative rarity of RAS-MAPK wild-type tumors and the relatively small sample sizes in these patient cohorts, this association need to be treated cautiously and more rigorously validated.

Rare germline variation in RAS pathway genes is known to impact nevus development, specifically strong activating germline mutations that cause developmental syndromes termed “RASopathies” [106]; patients with select RASopathies have higher number of melanocytic nevi [107–109]. Together with recent data potentially implicating other RAS-pathway members in melanoma/nevus risk loci, our data contribute to emerging evidence that common germline variation in genes regulating RAS pathway activity may also play a prominent role in melanoma risk and nevus development. We previously used a GWAS region-focused capture-HiC assay to nominate potential target genes from other melanoma GWAS loci [22], with *CBL* and *SHC1* nominated at separate loci. *CBL* encodes a negative regulator of receptor tyrosine kinases (and in turn, RAS signaling). We previously validated that the risk allele at the respective locus was associated with decreased *CBL* expression and demonstrated regulation of *CBL* by the region of the risk signal [22]. *SHC1*, which was nominated by the observation of active enhancer-promoter interactions from regions surrounding fine-mapped risk variants, encodes an adaptor protein that transduces signals from activated receptors to RAS by recruiting GRB2-SOS complexes, thereby promoting RAS activation and downstream RAS-MAPK signaling [110, 111]. In addition, the gene encoding another RAP GEF (*RAPGEF5*) is located at a melanoma risk locus identified by Landi and colleagues (rs12539524) [1], raising the possibility that other such GEFs could play a similar role. Lastly, a new GWAS of nevus count that nominated *NRAS* itself as a potential causal gene at a novel locus marked by rs7537232 [112]. While nearly every nevus acquires an oncogenic mutation in either *BRAF* or *NRAS*, our data suggest that germline variation regulating RAS activity may confer a growth or survival advantage prior to acquisition of a strongly activating somatic mutation.

We note several limitations of our study. Firstly, we have not positively identified the causal variant or variants underlying risk at this locus. Efforts to do so statistically are hampered by partial LD between QTL and melanoma risk signals, as well as a very large number of potential causal variants for melanocyte *RAPGEF1* expression. Nonetheless, the evidence implicating *RAPGEF1*, and perhaps *UCK1*, at this locus is strong. As noted above, we cannot rule out a role for *UCK1* in melanoma risk, further work will be required to assess potential phenotypes associated with variation in *UCK1* levels in melanocytes. Critically, while we have established that RAPGEF1 dysregulation leads to RAS activation, we have not demonstrated that RAPGEF1 acts as a direct GEF for RAS. We show a strong association of *RAPGEF1* levels with mutation status in melanoma tumors, however further work is required to establish whether and or to what degree RAPGEF1 actively contributes to RAS-MAPK pathway activation in tumors. Finally, the melanoma survival datasets are each relatively small, with even smaller numbers of RAS-MAPK pathway wild-type tumors; further assessment of associations of *RAPGEF1* with survival in patients with wild-type tumors is needed.

## Acknowledgements

This research was supported in part by the Intramural Research Program of the National Institutes of Health (NIH). The contributions of the NIH author(s) are considered Works of the United States Government. The findings and conclusions presented in this paper are those of the author(s) and do not necessarily reflect the views of the NIH or the U.S. Department of Health and Human Services. This work utilized the Biowulf cluster computing system at the NIH. The results appearing here are in part based on data generated by the TCGA Research Network. We would like to thank members at the National Cancer Institute Cancer Genomics Research Laboratory (CGR) for their help with sequencing efforts. We also thank all the cohorts, funders, and investigators who contributed to the melanoma GWAS, as originally acknowledged by Landi and colleagues. We would like to thank the research participants and employees of 23andMe. Mark Iles is supported in part by the National Institute for Health and Care Research (NIHR) Leeds Biomedical Research Centre. The views expressed are those of the author(s) and not necessarily those of the NHS, the NIHR or the Department of Health and Social Care. The Leeds Melanoma Cohort was funded by CRUK grants C588/A19167, C8216/A6129, C588/A10721 and NIH grant CA83115. JN is supported by Horizon Europe grant 101136622.

## Web Resources

- NIH Biowulf Cluster, http://hpc.nih.gov
- PLINK: https://www.cog-genomics.org/plink/
- GCTA-COJO: https://yanglab.westlake.edu.cn/software/gcta/
- RSparsePro: https://github.com/zhwm/RSparsePro_LD
- SuSiE: https: https://stephenslab.github.io/susieR/reference/susie_rss.html
- DAP-G: https://github.com/xqwen/dap
- AREsite2: http://nibiru.tbi.univie.ac.at/AREsite2/welcome
- Ensembl variant Effect Predictor: https://useast.ensembl.org/info/docs/tools/vep/index.html
- TWAS FUSION: http://gusevlab.org/projects/fusion/
- COLOC: https://github.com/chr1swallace/coloc
- SharePro: https://github.com/zhwm/SharePro_coloc
- ENCODE: https://www.encodeproject.org/encyclopedia/
- CHiCAGO: https://www.functionalgenecontrol.group/chicago
- HiCUP: https://www.bioinformatics.babraham.ac.uk/projects/hicup/
- LDlink: https://ldlink.nih.gov/?tab=home
- UCSC browser: https://genome.ucsc.edu
- WashU Epigenome Browser: http://epigenomegateway.wustl.edu
- Bedtools: https://bedtools.readthedocs.io/en/latest/
- Bedtoolsr: http://phanstiel-lab.med.unc.edu/bedtoolsr.html
- ROADMAP epigenomic project: https://egg2.wustl.edu/roadmap/web_portal/chr_state_learning.html
- TCGA melanoma cohort: https://www.cbioportal.org/study/summary?id=skcm_tcga_pan_can_atlas_2018
- Van Allen cohort data: https://www.cbioportal.org/study/summary?id=skcm_dfci_2015)
- Liu Cohort data: https://www.cbioportal.org/study/summary?id=mel_dfci_2019)
- Riaz cohort data: https://github.com/riazn/bms038_analysis/tree/master/data

## Data and Code Availability

Small RNA-seq data generated from 106 melanocyte cultures is currently being deposited on ArrayExpress. Melanocyte ATAC-seq and capture-HiC data are publicly available (ArrayExpress accession: E-MTAB-15079, E-MTAB-15048). Melanocyte genotyping, RNA-sequencing, and methylation data are previously published [13, 16] and accessible through NCBI database of Genotypes and Phenotypes (dbGaP) under accession dbGaP: phs001500.v2.p1.

MPRA are previously published [23]; raw sequencing can be accessed through NCBI Gene Expression Omnibus (GEO; https://www.ncbi.nlm.nih.gov/geo/) under the accession GEO: <u>GSE210356</u>.

Summary data for 2020 melanoma GWAS were obtained from dbGaP (dbGaP: phs001868.v1.p1), with the exclusion of self-reported data from 23andMe, Inc. and UK Biobank. The full GWAS summary statistics for the 23andMe discovery dataset will be made available through 23andMe to qualified researchers under an agreement with 23andMe that protects the privacy of the 23andMe participants. Please visit https://research.23andme.com/collaborate/#dataset-access/ for more information and to apply to access the data. Summary data from the remaining self-reported cases are available from the corresponding authors of that manuscript (Matthew Law; Mark Iles; and Maria Teresa Landi).

Melanoma tumor data for TCGA melanoma cohort was accessed using cBioportal database https://www.cbioportal.org/study/summary?id=skcm_tcga_pan_can_atlas_2018.

Leeds Melanoma cohort gene expression and mutation profiling dataset is available on European Genome-phenome Archive (study ID: EGAS00001002922, dataset ID: EGAD00010001561, dataset ID: EGAD00001008360). Lund Melanoma gene expression dataset is available on NCBI Gene Expression Omnibus (GSE65904); mutation profiling data was available from Cirenajwis *et al*., Supplementary Table 3 [72]. Liu and Van Allen cohort dataset was accessed using cBioportal (Liu cohort: https://www.cbioportal.org/study/summary?id=mel_dfci_2019) (Van Allen cohort: https://www.cbioportal.org/study/summary?id=skcm_dfci_2015) Riaz cohort mutation and expression dataset was accessed using https://github.com/riazn/bms038_analysis/tree/master/data

## Author Contributions

R.T., M.X, H.S., J.Yon, S.A.-S., M.L., T.R., H.S., G.D., L.J., T.M, T.Z., R.C., E.L., K.F., J.Yin, R.H., T.Z., G.J., D.T.B., J.N.-B., J.N., M.M.I., M.T.L., M.H.L., T.A., J.C., J.S., and K.M.B. contributed to the research activities described in this manuscript. K.M.B. led and supervised the research described. M.M. provided additional supervision and mentorship. R.T., M.X., and K.M.B. wrote the manuscript.

