## Supplementary Material for "Functional characterization of the 9q34.13 locus identifies *RAPGEF1* as modulating risk for melanoma and nevi via RAS activation"

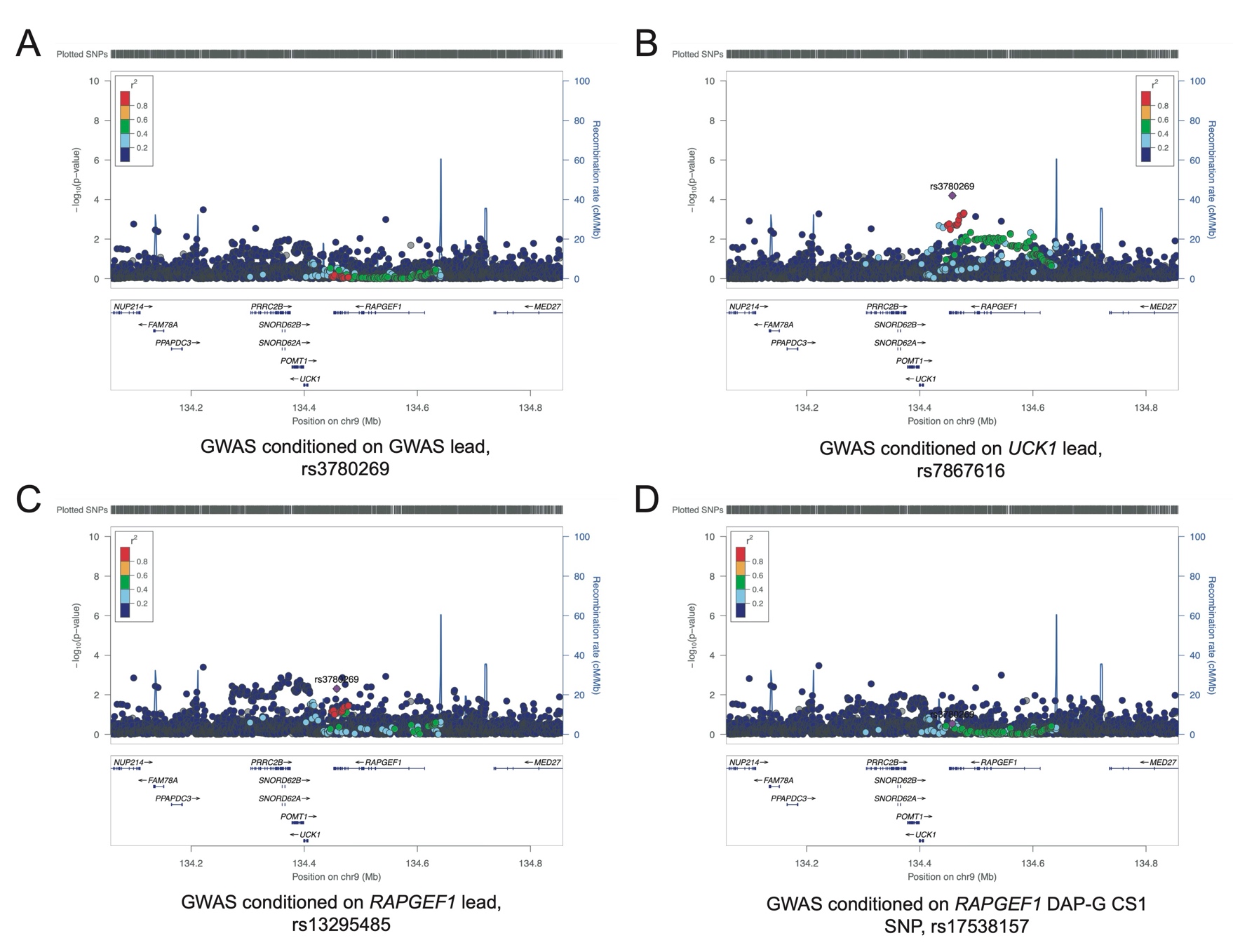

**Figure S1. Conditional analysis of the melanoma GWAS signal at a locus on chromosome band 9q34.13.** Manhattan plots of conditional melanoma GWAS data generated using the GCTA-COJO module. **A.** Melanoma GWAS conditioned on the lead GWAS variant (rs3780269); B. melanoma GWAS conditioned on the lead melanocyte *UCK1* eQTL variant (rs7867616); C. melanoma GWAS conditioned on the lead melanocyte RAPGEF1 eQTL variant (rs13295485); and D. melanoma GWAS conditioned on the highest posterior inclusion probability (PIP) SNP fine mapped by the Bayesian method DAP-G (rs17538157). Plots are colored to reflect linkage disequilibrium relative to the lead melanoma GWAS SNP (rs3780269).

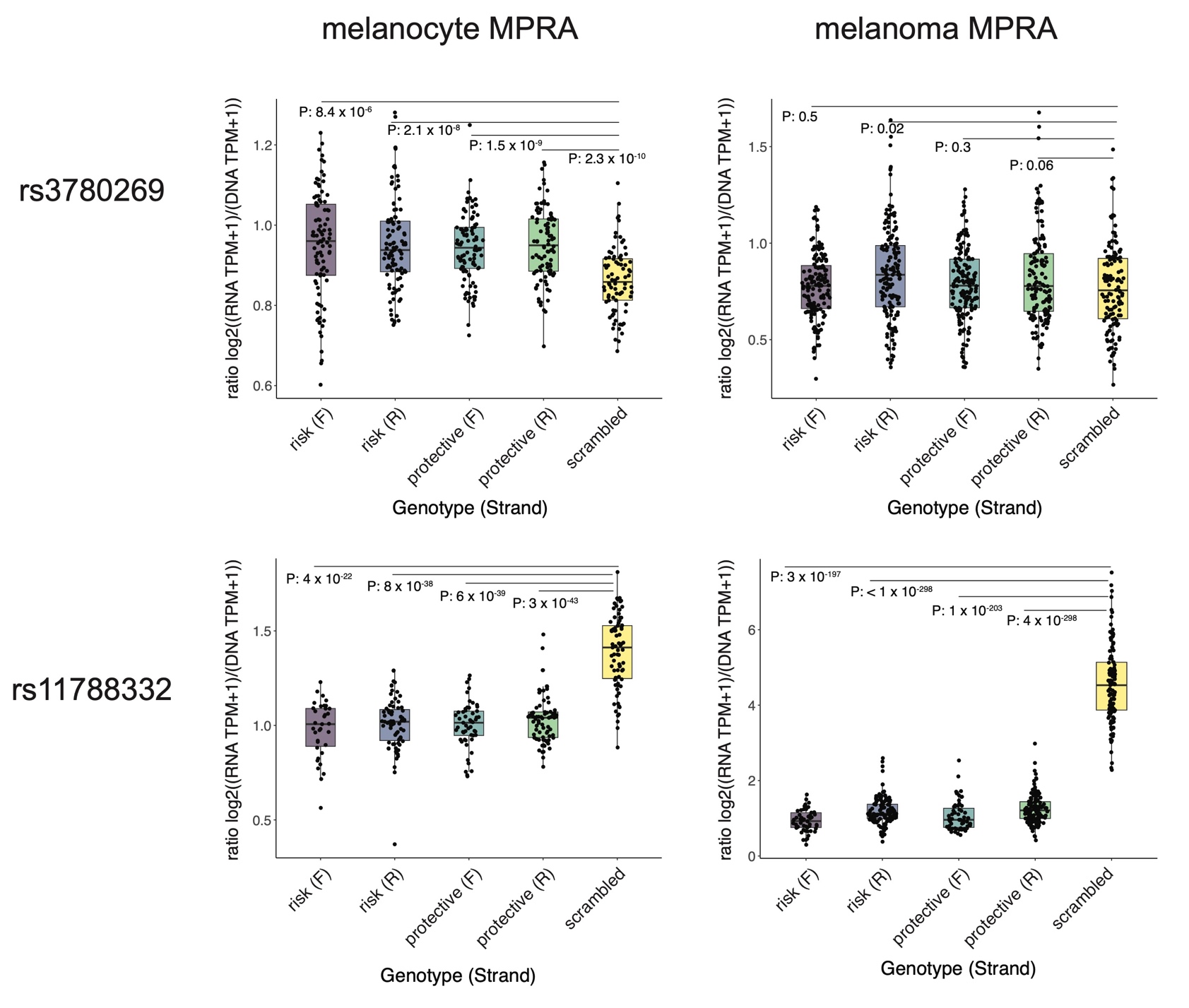

**Figure S2. Fine-mapped melanoma risk variants are located in regions with melanocyte and melanoma regulatory potential.** Two representative SNPs within regions demonstrating either significantly higher (rs3780269) or lower (rs11788332) regulatory activity relative to scrambled control sequences as assessed by Long and colleagues. The log2 ratio of RNA tags-per-million (TPM) relative to the DNA tags-per-million is shown for sequences harboring the melanoma risk or protective alleles cloned in forward and reverse orientations.

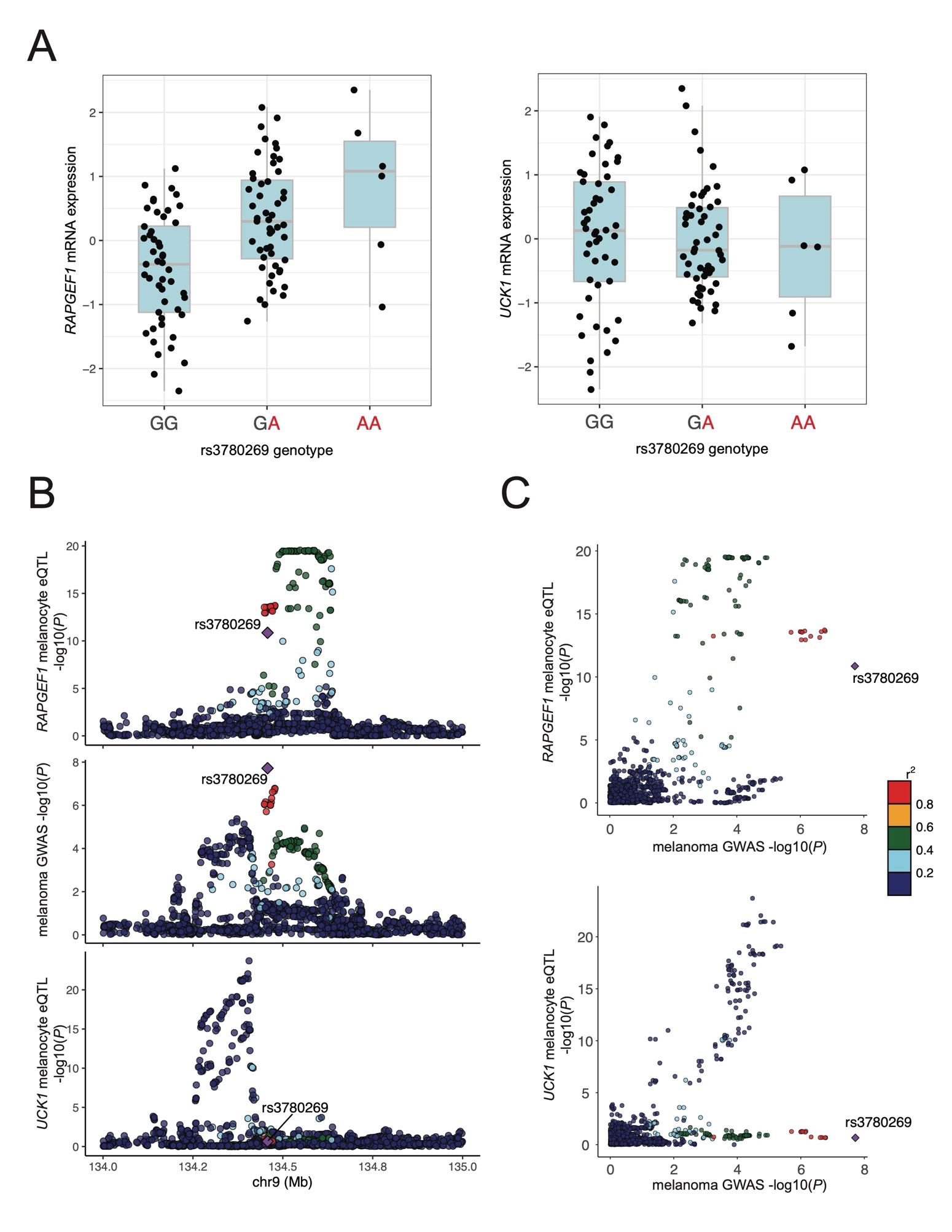

**Figure S3. Human primary melanocyte eQTLs for the lead melanoma risk SNP (rs3780269) and colocalization of risk and eQTL signals.** **A**. Box plots showing expression of *RAPGEF1* and *UCK1* in melanocytes homozygous or heterozygous for melanoma risk protective or risk alleles (rs3780269; risk allele is shown in red). **B**. Stacked Manhattan plots of data for the melanocyte *RAPGEF1* eQTL, the melanoma GWAS, and the melanocyte *UCK1* eQTL. **C**. LocusCompare colocalization plots comparing -log10 P-values for the melanoma GWAS to *RAPGEF1* and *UCK1* melanocyte eQTLs, respectively. Linkage disequilibrium for panels B and C is color coded relative to the lead melanoma GWAS SNP, rs3780269.

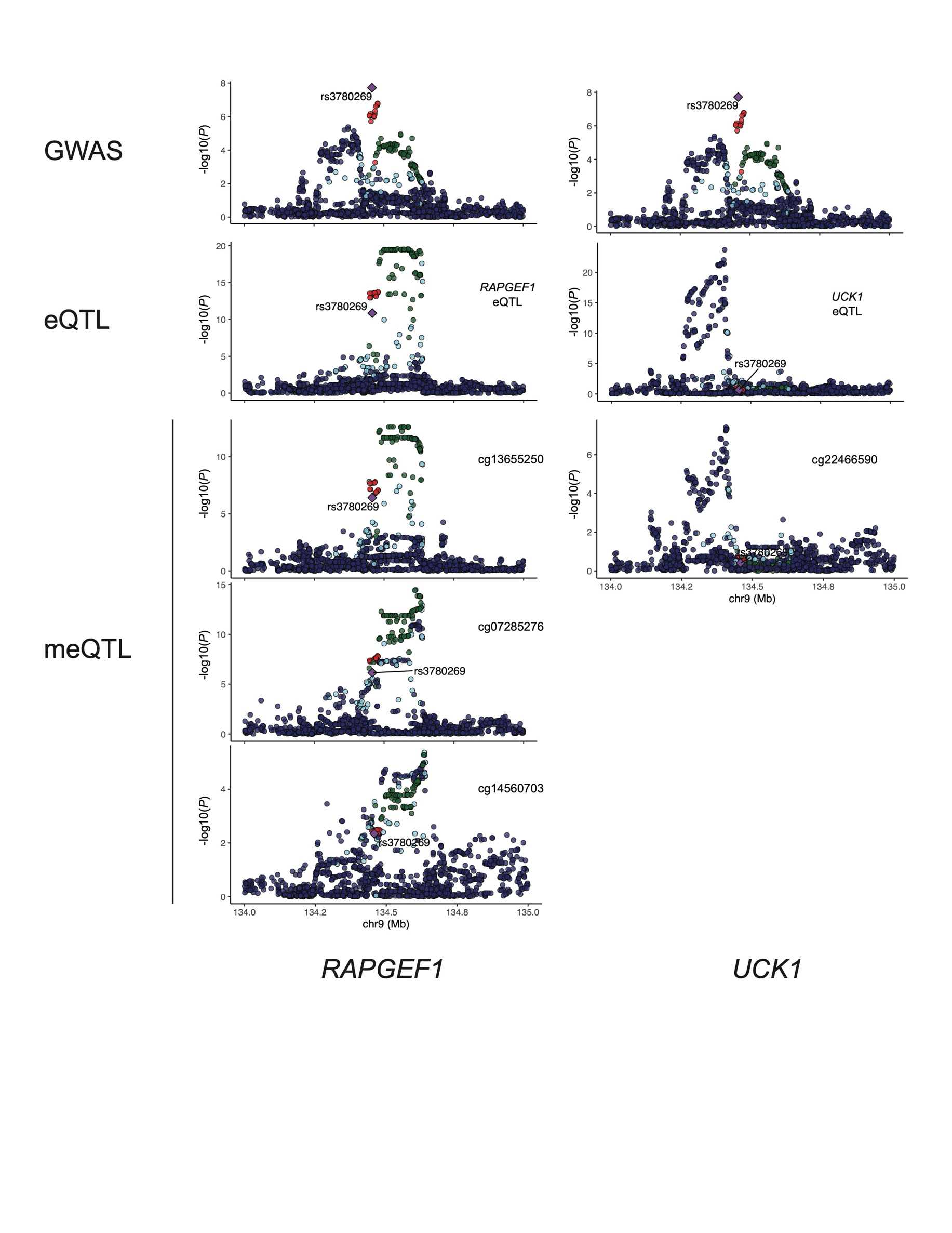

**Figure S4. Colocalization of melanoma risk with melanocyte eQTL and meQTL signals.** Stacked Manhattan plots of data for the melanoma GWAS, melanocyte *RAPGEF1* (left) or *UCK1* (right) eQTLs, and melanocyte meQTLs for CpG probes in *RAPGEF1* (left) or *UCK1* (right), respectively. Linkage disequilibrium is color coded relative to the lead melanoma GWAS SNP, rs3780269.
