## Supplementary Figures for "Functional characterization of the 9q34.13 locus identifies *RAPGEF1* as modulating risk for melanoma and nevi via RAS activation"

Rohit Thakur *et al.*

**Supplementary Methods**

*Tumor cohort descriptions*

*TCGA Melanoma Cohort (****Supplementary Table 2****)*: We assessed the extended dataset from the Cancer Genome Atlas (TCGA) skin cutaneous melanoma study [1], comprising 439 patient samples with matching mutation and expression profiling data accessed via the cBioPortal database [2-4], Skin Cutaneous Melanoma (TCGA, PanCancer Atlas) dataset (<https://www.cbioportal.org/study/summary?id=skcm_tcga_pan_can_atlas_2018>). This extended dataset was included as part of the PanCancer Atlas analyses [5, 6]. Mutation profiling was performed using whole-exome sequencing, and expression profiling was performed using RNA-sequencing. Four tumor purity measures were available [7] (PMID: 26634437). Only a small subset of samples in this dataset are primary tumors (n=76), whereas the majority represented metastatic tumors. In our analyses, we included only samples labeled as metastases (n=363); samples annotated as primary tumors were excluded. Among the remaining samples, 21 patients received ipilimumab, 2 patients received pembrolizumab, 1 patient received both ipilimumab and pembrolizumab, and 1 patient received nivolumab; mutation and gene expression profiles from the samples from these patients were generated from tumor samples collected prior to immunotherapy treatment. The vast majority of other samples were collected prior to treatment (treatments across these samples were varied), however a few (n=13) were collected following some form of treatment. Age at diagnosis, sex, tumor site-based staging, and all four estimates of tumor purity were used as covariates.

*Leeds Melanoma Cohort (****Supplementary Table 3,4****)*: We analyzed data from the population-based Leeds Melanoma Cohort [8, 9]. In this study, 2,184 patients were recruited between 2000 and 2012 from the northern United Kingdom; a subset of primary tumors from these patients collected within 3-6 months after initial diagnosis and prior to treatment were assessed for gene expression and somatic mutations. We only included samples where both transcriptome and mutation profiling data were available; we also excluded acral and mucosal melanomas, as well as melanomas arising at rare sites (such as vulva, vagina, anus, or penis that are not classified as mucosal in origin). In total, we analyzed tumors from 503 patients. Gene expression profiling was performed using the Illumina DASL platform [9] (European Genome-Phenome Archive, accession: EGAS00001002922); the probe ILMN-1769412, which targets all *RAPGEF1* transcripts, showed high-quality detection (signal above 95% of samples) and was selected as the representative probe for evaluating *RAPGEF1* expression in this cohort. All 503 were also profiled for mutations in *BRAF* (codons V600 and K601) and *NRAS* (codons Q61, G12, and G13). In a subset of patients (n=269), targeted sequencing of a panel of cancer-associated and melanoma-development-related genes (554 genes) was also performed [10] (European Genome-Phenome Archive, accession: EGAD00001008360). This gene panel included 35 of the 44 genes from the RAS-MAPK gene set described below. For the Leeds cohort, only a single tumor purity estimate was available, which was adjusted for in the analyses. For survival analyses, stage I tumors were excluded due to their high survival rate [11] (93-97% survival) and to better match other melanoma cohorts, which predominantly included more advanced-stage tumors. Age at diagnosis, sex, tumor purity, and AJCC staging were used as covariates.

*Lund Melanoma Cohort (****Supplementary Table 5****)*: We analyzed data from another previously published population-based melanoma cohort comprising tumor samples (n = 216) collected at the Department of Surgery, Skane University Hospital, between 2000 and 2012 [12, 13]. The cohort included a small proportion of primary tumors (n = 16) and predominantly metastatic samples (n = 188). As described previously, 95 patients (59%) did not receive treatment, while 67 patients (41%) underwent some other form of therapy [13]. Among those treated, 13 received neoadjuvant therapy (9 chemotherapy and 4 immunotherapy); targeted therapy was administered to 7 patients (4 with a BRAF inhibitor, 2 with imatinib, and 1 with sorafenib), and 2 patients received a vaccine. Of these, 146 patients had both gene expression and mutation data available and were subsequently analyzed. Gene expression profiling was performed using the Illumina Human-HT12v4.0 BeadChip platform (NCBI Gene Expression Omnibus, accession: GSE65904). In addition, targeted deep sequencing of 1,697 cancer-associated genes was conducted [14]; this gene panel included 32 of the 44 genes from the RAS-MAPK gene set (described below). Tumor purity was estimated using the ESTIMATE algorithm [15] implemented in the tidyestimate R-package (https://github.com/KaiAragaki/tidyestimate). This method infers tumor purity based on stromal and immune gene expression-based signatures. Age at diagnosis, sex, tumor purity, and AJCC staging were used as covariates.

*Liu Cohort (****Supplementary Table 6****)*: This cohort consisted of advanced-stage metastatic melanoma tumor samples [16] with matched gene expression and mutation profiling data (n = 121) available from the cBioPortal database (<https://www.cbioportal.org/study/summary?id=mel_dfci_2019>). All 121 patients in the study cohort received some form of immunotherapy treatment. For 116 of the 121 patients, tumor samples were retrospectively collected before anti-PD1 treatment (nivolumab and pembrolizumab). 17 out of 121 samples had previously received MAPK inhibition therapy, and 47 had been treated with anti-CTLA4 therapy (ipilimumab). In our analysis, we included samples annotated for “biopsy context” as “Pre-Ipi” or “Pre-PD1” and with “primary type” labelled as “skin”. We excluded all other primary types (acral, mucosal, and occult) and excluded all samples annotated either as “On-Ipi” or “On-PD1”. Gene expression profiles were generated by whole-transcriptome sequencing, and mutation data were derived from whole-exome sequencing. Applying these criteria, a total of 84 samples with both gene expression and mutation data were retained for downstream analysis. Tumor purity was estimated using the ESTIMATE algorithm and included as a covariate together with the purity variable provided in the publicly available dataset. Because age information was unavailable, the COSMIC aging signature-based value that was available was used as a surrogate covariate for age. Sex, AJCC stage, purity, purity estimate from ESTIMATE, and COSMIC aging signature were used as covariates.

*Van Allen Cohort (****Supplementary Table 7****)*: This cohort consists of patients with AJCC stage IV melanoma [17], for whom matched mutation and gene expression data (n=40) were available through the cBioPortal database (<https://www.cbioportal.org/study/summary?id=skcm_dfci_2015>). All patients in this cohort received immunotherapy (ipilimumab). Among them, 7 had previously undergone RAF inhibitor therapy, and 33 had received more than one prior treatment. Tumor samples were collected at a timepoint before the start of immunotherapy, and both expression and mutation profiling were done on these tumors. Gene expression data were generated using transcriptome sequencing, and mutational data were generated through whole-exome sequencing. Among the 40 patient samples with matching expression and mutation data, 35 originated from cutaneous (skin) melanoma primary tumor, 3 were occult, and 2 were mucosal. Owing to the relatively small size of this cohort compared with others described above, occult melanoma samples were retained due to their genetic similarity to cutaneous melanoma [18], whereas mucosal melanoma samples were excluded. Consequently, 38 tumor samples with both gene expression and mutation data were analyzed. Tumor purity estimate for the Van Allen cohort was available from the cBioPortal database. Since all patients were stage IV, the M staging information was used, with age at diagnosis, sex, and tumor purity also included as covariates.

*Riaz Cohort (****Supplementary Table 8****)*: Here, tumor samples were obtained from 68 metastatic melanoma patients [19], with tumor samples collected both before and during treatment with Nivolumab. Of the 68 patients, 33 had previously received ipilimumab. Whole-exome sequencing was performed to assess mutational profiles, and RNA-sequencing was used for gene expression. Because expression profiling data were available for matched pre-treatment and on-treatment samples, our analysis focused exclusively on data from pre-treatment samples (n=39). The gene expression data and mutation data were downloaded from <https://github.com/riazn/bms038_analysis/tree/master/data>. The downloaded gene expression counts were converted to TPM values using counts_to_tpm function (<https://github.com/skimlab/CCSBUtils>). Tumor purity measures were not available in the original meta-data; therefore, we estimated purity using the ESTIMATE algorithm implementation in the tidyestimate R-package. Because age and sex information were not available, only M staging and purity was used as a covariate.

*Tumor driver mutation definitions*

The driver annotations for tumor alterations were available on the cBioportal database for TCGA melanoma, as well as the Liu and Van Allen cohorts, which utilizes information from OncoKB [20, 21] and Cancer HotSpots [22, 23] databases for annotating an alteration as driver. Where available, gene fusion events classified as drivers were included in the mutant subgroup classifications. For the remaining three other cohorts, the alterations were defined as driver mutations for oncogenes via manual look up on the OncoKB and Cancer HotSpots databases. For tumor suppressor genes such as *NF1* across these cohorts, we used the ANNOVAR [24] tool for annotating variant predicted effect across 11 prediction algorithms, SIFT [25], PolyPhen2 [26], Mutation Assessor [27], SVM radial, LR predict, MetaSVM and MetaLR [28], PROVEAN [29], PrimateAI [30], AlphaMissense [31], and CLNSIG [32]. An alteration was considered as a potential driver if predicted pathogenic or likely pathogenic by the majority of these tools. This integrative approach combined expert curated driver annotations with computational predictions to provide comprehensive information for classification of gene alterations.

**Supplementary Results**

*Melanocyte splice QTL (sQTLs)*

Splice QTL (sQTL) analysis of primary melanocytes [33] did not identify FDR-significant findings for the lead melanoma GWAS SNP rs3780269; we observe only a weak association for increased usage of the chr9:134477536-134497183 junction (hg19, *P* = 0.04) and decreased usage of chr9:134479440-134497183 junction (*P* = 0.06) of *RAPGEF1*, as well as decreased usage of the chr9:134404588:134404875 *UCK1* junction (*P* = 0.05) with the risk-allele. These sQTLs appear to be merely overlapping with the melanoma risk signal, driven by independent SNPs with low LD to the melanoma risk signal (lead rs58411788, *P* = 7.3 x 10^-5^, r^2^ = 0.002, D’ = 0.12, 1000G EUR; rs142737053; *P* = 2.2 x 10^-4^, r^2^=0.0007, D’=0.13; and rs142737053, *P* = 6.0 x 10^-5^, r^2^=0.0007, D’=0.13, respectively), and we observe no statistical evidence of colocalization.

*Micro-RNA binding analysis for fine-mapped variant rs3739497*

One fine-mapped melanoma risk variant, rs3739497 (*P*_GWAS_ = 1.9 × 10⁻⁶; OR = 0.95; risk allele = A; protective allele = G; r² to GWAS lead = 0.88), is located in the 3′ untranslated region (UTR) of *RAPGEF1*, raising the possibility that it could affect post-transcriptional regulation of *RAPGEF1* by altering a microRNA binding site or an AU-rich element (ARE) motif [34]. While we observed no overlap with ARE motifs, we did identify predicted binding sites for *miR-124-3p* and *miR-506-3p* within 50 bases of the variant. Notably, Ramachandran and colleagues previously tested for an allele specific effect for this variant on regulation of transcript stability by *miR-124-3p* using a reporter system in HEK293 cells and observed a stronger reduction in transcript levels with the melanoma risk allele of rs3739497 relative to the protective allele [35]. We observe *miR-124-3p* to be present only at very low levels in small RNA sequencing data from a panel of 106 human primary melanocytes (expressed in the 1^st^ percentile of all small RNA transcripts) as well as melanoma tumors from TCGA (expressed in only 58 of 448 samples assayed for both miRNA and mRNA) [1]. We also observe no correlation between *miR-124-3p* and *RAPGEF1* levels across melanocyte cultures (Spearman’s ρ = 0.06; *P* = 0.49). Of the TCGA melanoma tumors with detectable *miR-124-3p* and *RAPGEF1* expression, we likewise observe little correlation (Spearman’s ρ = 0.19; *P* = 0.1). Furthermore, the allelic direction of effect of *miR-124-3p* on transcript stability observed by Ramachandran and colleagues is opposite of the direction of allelic *RAPGEF1* expression observed in primary melanocytes (described below), making regulation by *miR-124-3p* an unlikely mechanism for the causal sequence variant at this locus. In contrast, *miR-506-3p* is indeed highly expressed in both cell types (93^rd^ percentile in melanocytes, expressed in 341/448 melanomas), however, *miR-506-3p* expression showed little correlation with *RAPGEF1* in either dataset (melanocyte Spearman’s ρ = -0.01, *P* = 0.89; melanoma Spearman’s ρ = 0.06, *P* = 0.25). Collectively, these data suggest rs3739497 is unlikely to drive allelic variation in *RAPGEF1* transcript levels via allelic micro-RNA binding; we nonetheless cannot rule out a potential translational effect.

*Melanocyte methylation QTLs (meQTLs)*

To find further support for *RAPGEF1*, *UCK1*, or other candidates, we also assessed allele-specific methylation. Specifically, we noted a pattern of allele-correlated CpG methylation near *RAPGEF1* and *UCK1* in methylation QTL (meQTL) data from primary melanocytes [36] similar to that observed for expression of both genes. Broadly, we observe numerous FDR-significant meQTLs across the larger region surrounding the melanoma association signal (**Supplementary Table 36**). Using a melanocyte methylation-wide association study (MWAS) approach to identify those meQTLs closely associated with the melanoma risk signal, we identify four CpG probes whose predicted methylation was significantly associated with melanoma risk (**Supplementary Table 37**). Three were annotated as located within *RAPGEF1* (cg13655250; *P*_TWAS_ = 5.90 x 10^-5^, low methylation->risk; cg14560703, *P*_TWAS_ = 3.99 x 10^-3^, low methylation->risk; cg07285276, *P*_TWAS_ = 6.86 x 10^-3^; high methylation->risk) and one within *UCK1* (cg22466590; *P*_TWAS_ = 7.48 x 10^-8^; high methylation->risk). At the single-signal level, the meQTL signals are strikingly similar to and colocalize with eQTL signals for the respective genes (**Supplementary Table 38, 39**; **Supplementary Figure 4**) but not the melanoma GWAS signal itself (**Supplementary Tables 40, 41**), reinforcing *RAPGEF1* and *UCK1* as strong causal candidates regulated by a complex pattern of allelic *cis*-regulatory variants in melanocytic cells.

4.	de Bruijn, I., R. Kundra, B. Mastrogiacomo, T.N. Tran, L. Sikina, T. Mazor, X. Li, A. Ochoa, G. Zhao, B. Lai, A. Abeshouse, D. Baiceanu, E. Ciftci, U. Dogrusoz, A. Dufilie, Z. Erkoc, E. Garcia Lara, Z. Fu, B. Gross, C. Haynes, A. Heath, D. Higgins, P. Jagannathan, K. Kalletla, P. Kumari, J. Lindsay, A. Lisman, B. Leenknegt, P. Lukasse, D. Madela, R. Madupuri, P. van Nierop, O. Plantalech, J. Quach, A.C. Resnick, S.Y.A. Rodenburg, B.A. Satravada, F. Schaeffer, R. Sheridan, J. Singh, R. Sirohi, S.O. Sumer, S. van Hagen, A. Wang, M. Wilson, H. Zhang, K. Zhu, N. Rusk, S. Brown, J.A. Lavery, K.S. Panageas, J.E. Rudolph, M.L. LeNoue-Newton, J.L. Warner, X. Guo, H. Hunter-Zinck, T.V. Yu, S. Pilai, C. Nichols, S.M. Gardos, J. Philip, A.P.G.C. Aacr Project Genie Bpc Core Team, K.L. Kehl, G.J. Riely, D. Schrag, J. Lee, M.V. Fiandalo, S.M. Sweeney, T.J. Pugh, C. Sander, E. Cerami, J. Gao, and N. Schultz, *Analysis and Visualization of Longitudinal Genomic and Clinical Data from the AACR Project GENIE Biopharma Collaborative in cBioPortal.* Cancer Res, 2023. **83**(23): p. 3861-3867.

26.	Adzhubei, I., D.M. Jordan, and S.R. Sunyaev, *Predicting functional effect of human missense mutations using PolyPhen-2.* Curr Protoc Hum Genet, 2013. **Chapter 7**: p. Unit7 20.
